# Zooarchaeological investigation of the Hoabinhian exploitation of reptiles and amphibians in Thailand and Cambodia with a focus on the Yellow-headed Tortoise (*Indotestudo elongata* (Blyth, 1854))

**DOI:** 10.1101/2023.04.27.538552

**Authors:** Corentin Bochaton, Sirikanya Chantasri, Melada Maneechote, Julien Claude, Christophe Griggo, Wilailuck Naksri, Hubert Forestier, Heng Sophady, Prasit Auertrakulvit, Jutinach Bowonsachoti, Valéry Zeitoun

## Abstract

While non-marine turtles are almost ubiquitous in the archaeological record of Southeast Asia, their zooarchaeological examination has been inadequately pursued within this tropical region. This gap in research hinders a complete comprehension of past human subsistence strategies and economies, as only a limited number of comprehensive studies encompassing all the taxa found in archaeological sites have been conducted thus far. This constraint becomes particularly significant in relation to prehistoric hunter-gatherer populations, who might have extensively utilized inland chelonian taxa. In order to initiate a new approach to the study of past human-turtle interactions in Southeast Asia, we propose an in-depth zooarchaeological analysis of turtle bone remains recovered from four Hoabinhian Hunter-gatherer archaeological assemblages located in Thailand and Cambodia, dating from the Late Pleistocene to the first half of the Holocene. Our study focuses on the bone remains attributed to the Yellow-headed Tortoise (*Indotestudo elongata*) as it is the most represented taxon in archaeological assemblages in the region of study. For this species, we developed osteometric equations enabling the estimation of the carapace size of the archaeological individuals. This allowed us to study the size structure of the archaeological populations at different sites and to reveal the human exploitation strategies of these animals. We observed a significant taphonomic homogeneity among the studied assemblages, along with similarities in the diversity of hunted reptile and amphibian taxa as well as the size of the exploited tortoises. These findings suggest consistent subsistence behaviors across distinct sites, despite their varying environmental conditions, and raise the possibility of cultural similarities across different periods and regions. Additionally, we provide a baseline for future zooarchaeological studies and a methodological framework for the detailed analysis of archaeological turtle bones in continental Southeast Asia.

## Introduction

The Hoabinhian has been a significant topic in prehistoric research in Mainland Southeast Asia for nearly 90 years. Since its first definition by the French archaeologist Madeleine Colani in the early 1930s (Collectif, 1932), the Hoabinhian has undoubtedly been one of the most debated subjects in the field. Various aspects related to the Hoabinhian populations have been discussed, including their spatial and temporal distribution, definition, technological and functional characteristics of their lithic industries, economic organization, and environmental context (Forestier et al., 2021).

The chronology of the Hoabinhian has significantly expanded since the discovery of the first characterized Hoabinhian sites. Currently, the earliest Hoabinhian deposits is considered to be the Xiaodong Rock shelter in Southwest China, dating back to about 43,000 BP (Ji et al., 2016).Similarly, the site of Huai Hin in Northwest Thailand, dated to approximately 3700 BP, is considered the last occurrence with a lithic production associated with ceramic sherds (Forestier et al., 2013; Zeitoun et al., 2008). Over a hundred Hoabinhian sites have been reported in Southeast Asia (Chung, 2008; Forestier et al., 2017; Moser, 2001; White, 2011; Zeitoun et al., 2008), raising questions about the variability of Hoabinhian lithic assemblages, including unexpected operational sequences on pebble matrix (Forestier et al., 2023, 2022). Nonetheless, despite these findings, the Hoabinhian people remain poorly understood from both cultural and material perspectives. The perceived homogeneity and limited diversity in their lithic material culture, potentially influenced by their extensive reliance on perishable vegetal resources (Forestier, 2003), pose a challenge in characterizing the activities undertaken at the sites, and in determining their overall functions (e.g., long-term occupation, butchering site, hunting camp). Additionally, the archaeological evidence paints an improbable image of cultural stagnation spanning over 30,000 years among diverse hunter-gatherer groups across an expansive region with varying environmental, ecological, and geographic conditions (Zeitoun et al., 2019).

White (2011) proposed that cultural diversity in mainland Southeast Asia began to emerge precisely in the late Late Pleistocene, which is also suggested by burial practices (Imdirakphol et al., 2017). Forestier et al. (2013) argued that further analyses were needed to evaluate the entire corpus of Southeast Asian lithic industries to describe potential “cultural variations”, and since then, certain patterns have started to emerge (Forestier et al., 2021, 2017). Nevertheless, the lack of detailed zooarchaeological analyses renders our knowledge of the Hoabinhian paleoecology and subsistence strategies somewhat unclear. This also hinders the characterization of archaeological deposits, as faunal data are of paramount importance in understanding the use and occupation periods of the sites.

The economic aspects of the Hoabinhian culture have been addressed by several authors (Glover, 1977; Gorman, 1971, 1970, 1969; Vu, 1994; Yen, 1977). However, pioneering prehistoric zooarchaeological studies relying on occurrence data (Gorman, 1971) only offer a limited understanding of the choices made by the hunters, as they focus on qualitative data and the diversity of the exploited animals rather than quantitative information reflecting the intensity of the exploitation of each species. These initial works also lack the detailed taphonomic and taxonomic analyses needed to describe past human behaviors and bone accumulation processes more comprehensively. As a result, we currently have only a vague idea of the potentially strong spatial and chronological variability of the subsistence strategies of the Hoabinhian people that have occupied and exploited a wide diversity of tropical environments across an extensive period of time. These issues in Southeast Asian zooarchaeology have been thoroughly reviewed by Conrad (2015). One of the identified issues is the lack of detailed study of each animal group, especially the non-mammal taxa, including reptiles and non-marine turtles, which often account for a significant portion of the animal bone assemblages found in the archaeological record.

This limitation is not unique to continental Southeast Asia but is more pronounced here than in many other areas. Zooarchaeological studies fully focused on non-marine turtles have been conducted in non-tropical regions such as Europe (Blasco, 2008; Nabais et al., 2019; Nabais and Zilhão, 2019), the Near East (Biton et al., 2017; Blasco et al., 2016; Speth and Tchernov, 2002), South Africa (Avery et al., 2004; Thompson and Henshilwood, 2014), and North America (Rhodin, 1992). However, such studies have been limited in tropical areas, including Southeast Asia, despite the fact that turtle bones are more commonly found in tropical settings than in temperate regions where prehistoric studies have a strong tradition. In Southeast Asia, this problem partly stems from the general lack of detailed anatomical data allowing for the identification of taxa based on isolated plate remains, and the absence of appropriate methodological frameworks to analyze this material. Several works have been conducted regarding the osteology of Southeast Asian turtles, most of which focusing on the Geoemydidae family to address questions related to phylogeny and paleo-biodiversity (Garbin et al., 2018; Naksri, 2013, 2007; Naksri et al., 2013). However, only a few works have focused on the study of isolated elements found in the archaeological record (Claude et al., 2019; Pritchard et al., 2009). Despite these limitations, some zooarchaeological studies of Hoabinhian archaeological deposits in mainland Southeast Asia have started to characterize the exploitation of non-marine turtles by prehistoric populations. In the few existing studies, turtle remains are often left unidentified: Ban Rai Rockshelter (Treerayapiwat, 2005); Banyan Valley Cave (Higham, 1977); Gua Gunung Runtuh (Zuraina, 1994); Gua Harimau (Bulbeck, 2003); Gue Kechil (Dunn, 1964; Medway, 1969); Gua Ngaum (Bulbeck, 2003); Gua Peraling (Adi, 2000); Gua Teluk Kelawar (Bulbeck, 2003); Tham Lod Rockshelter (Amphansri, 2011); Lang Kamnan Cave (Shoocongdej, 1996); Khao Toh Chong Rockshelter (Van Vlack, 2014); Moh Khiew II Rockshelter (Auetrakulvit, 2004); Spirit Cave (Higham, 1977); Tham Phaa Can (Higham, 2002); Thung Nong Nien Rockshelter (Auetrakulvit, 2004). In some studies, they are identified but not quantified by species, for instance, in the Lang Rongrien Rock Shelter assemblage (Anderson, 1990; Mudar and Anderson, 2007). In the rare studies in which turtle bones are identified and quantified, the Yellow-headed Tortoise (*Indotestudo elongata* (Blyth, 1854)) often stands out as the most abundant species, at least in its modern distribution area, in sites such as Doi Pha Kan Rockshelter (Frère et al., 2018), Laang Spean Cave (Frère et al., 2018), and Spirit Cave (Conrad et al., 2016). Outside of the modern range of that species, Geoemydidae turtles are best represented in Malaysian sites such as Gua Sagu (Rabett, 2012) and Gua Tenggek (Rabett, 2012). There is also a site in Vietnam (Hiem Cave) where the tortoise *Manouria* is the most abundant turtle taxon, but the small size of the faunal assemblage does not allow drawing conclusions from this observation (Masojc et al., 2023). However, even in the quantified studies mentioned above, the study of the turtle bone remains is still superficial. For instance, a detailed analysis of the taphonomy of the bone assemblages is not conducted, and the population of turtles exploited is not characterized beyond the aspect of its species composition.

To address these issues and to provide the first detailed data regarding the prehistoric exploitation of Southeast Asian turtles, we conducted an in-depth zooarchaeological analysis of turtle bone remains recovered from four hunter-gatherer archaeological assemblages located in Thailand and Cambodia, dating from the Late Pleistocene to the first half of the Holocene. These sites are the Doi Pha Kan rockshelter, the Moh Khiew Cave, and the Khao Tha Phlai Cave in Thailand, and the Laang Spean Cave in Cambodia. Additionally, to gain a more precise understanding of the exploitation strategies of non-marine turtles by archaeological human populations, we developed osteometric equations enabling the estimation of carapace size for the archaeological individuals of *Indotestudo elongata*. We chose to focus on this species as most of the rich assemblages of turtle bones collected in the four considered sites correspond to this species (Frère *et al*. 2018; C. B., J. C. preliminary observations). This analytical tool allows us to study the size structures of the archaeological populations at different sites and to characterize the choices made by the hunters. Together, these data provide the first characterization of the exploitation of non-marine turtles by Pleistocene and Holocene hunter-gatherer populations of continental Southeast Asia.

## Material and Methods

### Main Characteristics of *Indotestudo elongata*, the Yellow-headed Tortoise

The genus *Indotestudo* currently includes three species: Forsten’s Tortoise *Indotestudo forstenii* (Schlegel & Müller, 1845), Travancore Tortoise *Indotestudo travancorica* (Boulenger, 1907), and Yellow-headed Tortoise *Indotestudo elongata* (Blyth, 1854), with the latter two being sister taxa (Iverson et al., 2001). These species are distributed in India (*I. travancorica*), Sulawesi (*I. forstenii*), and northern India and continental Southeast Asia (*I. elongata*) (Rhodin et al., 2021). *Indotestudo elongata* is the only species of the genus present in continental Southeast Asia and is found in most areas of Thailand, Cambodia, Vietnam, and Laos (Ihlow et al., 2016; Rhodin et al., 2021). It is also present in northwestern Malaysia but is absent from most of this country and the insular Sunda (Ihlow et al., 2016).

*Indotestudo elongata* is a medium-sized tortoise, with adult individuals reaching a Straight Carapace Length (SCL) of about 300 mm (Taylor, 1970), while the largest recorded specimen was a male with an SCL of approximately 380 mm (Rhodin et al., 2021). Sexual dimorphism may vary among populations, but males are generally larger than females (Ihlow et al., 2016). Hatchlings typically measure between 50 to 55 mm SCL (Ihlow and Handschuh, 2011). The species is found in various environments, including different types of forests (Ihlow et al., 2016). *I. elongata* is active during the daytime, particularly in the early morning and late afternoon. It displays seasonal activity patterns, aestivating during the dry season (from March to May) to avoid the highest temperatures, and spends much of its time hiding in former burrows of other animal species, including porcupines (Ihlow et al., 2016; Som and Cottet, 2016; van Dijk, 1998). During the rainy season (from July to October), individuals spend most of their time in open areas, moving to more closed environments, mostly semi-evergreen and pine forests, when the climate becomes drier (Som and Cottet, 2016; van Dijk, 1998). Depending on their sex, individuals reach sexual maturity between 175 mm and 240 mm SCL at an age of 6-8 years old (Eberling, 2011; Sriprateep et al., 2013; van Dijk, 1998), and reproduction typically takes place during the rainy season.

### Presentation of the Studied Sites and Assemblages

#### The Doi Pha Kan rockshelter

The Doi Pha Kan rockshelter is located in northern Thailand (E 99° 46‘ 37.2‘‘ ; N 18° 26‘ 57.0‘‘). It has been under study since 2011 by P. Auetrakulvit and V. Zeitoun, and its excavation is still ongoing. The site is well-known for its three Hoabinhian burials, which have been dated between 11,170 ± 40 and 12,930 ± 50 BP (Imdirakphol et al., 2017; Zeitoun et al., 2019). Two of these sepultures contained turtle shell elements in anatomical connection, interpreted as funeral offerings. The site has also provided a rich archaeological assemblage corresponding to a Hoabinhian occupation older than the sepultures. The sedimentary stratigraphy of the site is relatively homogeneous, and as a result, its archaeological material has been considered as a single assemblage thus far.

Samples of the lithic material and animal bone assemblages collected from the site have already been the object of studies (Celiberti et al., 2018; Frère et al., 2018), but much of the material is still under study. In this paper, we will present zooarchaeological data collected on the herpetofaunal taxa bone remains recovered at the site until 2019. To collect this dataset, the entire archaeofaunal material has been observed to extract and study the reptile and amphibian bone remains. The material screened in this way corresponds to the material previously studied by S. Frère (Frère et al., 2018), with the addition of the material collected following this first study. In the initial study, S. Frère analyzed 4256 animal remains, of which 2541 were attributed to vertebrate species. So far, no complete study of this faunal assemblage has been completed, and no Minimum Number of Individuals (MNI) data have been published. In these conditions, the overall weight of the herpetofauna we collected in respect to the full sample cannot be assessed with precision. However, the existing data indicates it could account for a significant portion of the full assemblage, as it corresponds to 17.1% of the total bone weight and 51% of the vertebrate total Number of Identified Specimens (NISP) analyzed in the previous study (Frère et al., 2018).

#### The Moh Khiew Cave

The site of Moh Khiew Cave is a 30m long archaeological rock-shelter located in southern Thailand, in the Krabi Province (E 98° 55’ 49.27’’; N 08° 09’ 36.32’’). It was first excavated by S. Pookajorn between 1991 and 1994 (Pookajorn, 2001) before being the object of a later excavation in 2008 by P. Auetrakulvit to clarify its stratigraphy (Auetrakulvit et al., 2012). The stratigraphy of the site is composed of several archaeological layers dated from the Holocene to the Late Pleistocene, most of which correspond to Hoabinhian occupations. During the excavations, five sepultures were also discovered: one during the first excavations and four in 2008.

Regarding the zooarchaeological data, the complete assemblage from the first excavations was studied by P. Auetrakulvit (Auetrakulvit, 2004). In this assemblage, the MNI data indicate that herpetofaunal species account for 24.9% of the assemblage. This proportion dramatically increases to more than 70% of the material if the NISP is considered, with nearly 60% attributed to non-marine turtle bone alone. Unfortunately, the turtle remains were not identified further at the time of this first study, and we were unable to locate this material for reanalysis in the present study. However, we had access to the material collected during the 2008 excavation, but only to previously studied herpetofaunal bone samples that were extracted from the complete bone assemblage of the excavation by several master students. These bones were recovered from the four different layers identified during the 2008 excavation of Moh Khiew Cave (Auetrakulvit et al., 2012). The first level (Layer 1) is composed of the upper 90 cm of the sequence and is a disturbed layer that has not been dated, corresponding to the levels 1 and 2 identified by S. Pookajorn in the first excavations. The second layer includes the sediment collected between 90 and 170 cm of depth and has been radiocarbon dated with three dates between 7520+-420 BP and 8660+-480 BP. This layer corresponds to level 3 identified in the previous excavations. The third layer corresponds to a depth between 170 and 210 cm and has been dated with two radiocarbon dates of 8730+-480 BP and 9270+-510 BP. It was also identified as corresponding to level 3 previously described by S. Pookajorn. The last layer corresponds to the remaining stratigraphy, a layer of scree, and was not dated but associated with level 4 described in the previous excavations.

#### The Khao Tha Phlai Cave

The site of Khao Tha Phlai is a cave located in southern Thailand, in the province of Chumphon (E 10°36’12.39“; N 99° 5’49.08”). It has been excavated by the 12^th^ Regional Office of Fine Arts Department, Nakhon Si Thammarat, since 2014, and so far, has been the object of two excavation campaigns with two test-pits conducted in the deposit. The first test-pit of 9 m² (TP1) was in 2014, and the second of 20 m² (TP2) was in 2021-2022.

The archaeofaunal material collected in TP1 has been the object of a preliminary study by S. Jeawkok in the framework of a Bachelor thesis, which remained unpublished. This first study conducted on 6945 bone remains indicated that reptiles represent around 25% of the NISP of the complete assemblage, with the remaining bones mostly attributed to large mammals. In the present study, we considered the herpetofaunal material previously extracted by S. Jeawkok from the faunal samples collected in TP1. We also consulted the full archaeofaunal sample recovered from TP2 to extract the reptile and amphibian bones from it. We considered the samples collected in the two test-pits separately and subdivided the samples into two assemblages corresponding to Metal Ages (between 75 and 130 cm of depth in TP2 and between 65 and 180 cm in TP1) and Neolithic periods (between 130 and 320 cm of depth in TP2 and between 180 and 320 in TP1). These layers have been dated based on the typology of the archaeological artifacts (lithic tools, ceramic shards, and metal objects) they have provided.

#### The Laang Spean Cave

The site of Laang Spean is a large cave of more than 1000 m² located in northwest Cambodia, in the Battambang province (E 102° 51‘ 00.0‘‘; N 12°51‘ 00.0‘‘). The site was first excavated between 1965 and 1968 by R. Mourer and C. Mourer-Chauviré (Mourer-Chauviré et al., 1970; Mourer-Chauviré and Mourer, 1970). The archaeofaunal material collected during these excavations has been the object of a preliminary study, but with the exception of the rhinoceros remains (Guerin and Mourer-Chauviré, 1969), no in-depth paleontological study has been conducted, and no zooarchaeological analysis was performed. Following this first exploration, the site was the object of a new detailed archaeological excavation by H. Forestier between 2009 and 2019, aiming to document the Hoabinhian occupation previously identified in more detail (Forestier et al., 2015; Sophady et al., 2016). These excavations, conducted on a surface of over 40 m² led to the discovery of an important undisturbed Hoabinhian layer dated between 5018 ± 29 cal. BP and 10,042 ± 43 cal. BP, as well as several Neolithic burials dug into the Hoabinhian level (Zeitoun et al., 2021, 2012). These sepultures have been dated from 3335 ± 30 to 2960 ± 30 BP (Sophady, 2016).

Regarding the subdivision of the material, the archaeological remains collected in the squares lacking traces of Neolithic perturbations (see Forestier et al., 2015) have been associated with the Hoabinhian occupation. There is no stratigraphic evidence suggesting a subdivision of this Hoabinhian assemblage, obviously representing several occupations over a time span of more than 5000 years. On the other hand, the squares in which sepultures were found are grouped together under the term “Sepulture layer”, and the first 120 cm of the disturbed squares are considered as being a “Neolithic layer”. However, these subdivisions are very artificial and probably correspond to a mix of Neolithic and Hoabinhian material, as evidenced by the lithic material from these contexts (H. Forestier, com. pers.).

A sample of the complete faunal assemblage collected in the Hoabinhian squares has been the object of a first zooarchaeological study by S. Frère (Frère et al., 2018). In this first study, among the 5885 vertebrate remains identified, turtles account for 44% of the NISP, monitor lizards for 1%, and large mammals for 37%. Unfortunately, no MNI data were reported, and the fact that most of the small fragments were not attributed to at least a size class of animal makes these results difficult to interpret. The study presented in this paper corresponds to the herpetofaunal material collected in the complete assemblage of bones recovered during all the excavations since 2009. This material has been extracted by C. B. upon the consultation of all the bone samples collected on the site. The zooarchaeological study of the other groups of vertebrate for the complete Hoabinhian assemblage of Laang Spean is currently in progress, and the final results are not available at the moment.

### Quantification of the Zooarchaeological data and Taphonomic Analyses

The basic units of quantification considered are the Number of Identified Specimens (NISP) and the Weight of the Remains (WR) (Lyman, 2008). The fragmentation of each bone has been recorded by describing the Percentage of Completion (PC) of the anatomical elements represented by the fragment. The anatomical side of the bones has been registered when possible for the best represented and easiest to identify anatomical elements (i.e., peripheral plates and all paired elements of the plastron). A Minimum Number of Elements (MNE) has been calculated for each anatomical part to assess differences in skeletal element representation across various archaeological contexts. To do so, we added the PCs of a given element and divided the result by 100. The results were rounded up to the nearest higher whole number to obtain the MNEs. The Minimum Number of Individuals (MNI) is defined using the anatomical element with the highest MNE (Lyman, 2008). The anatomical distributions are represented by the Percentage of Representation (PR) of Dodson and Wexlar (1979) using the MNE of each anatomical element and the MNI of the considered assemblage. All the archaeological bones have been weighed individually. To avoid potential impact of taxonomic identification bias on the anatomical distribution of the remains, we considered all the turtle/tortoise taxa in these analyses and not only the bone fragments attributed to *Indotestudo*. As tortoises are the best-represented taxa in the different assemblages with consistently more than 60% of the NISP attributed to that group, most of these unidentified turtle bones likely represent *Indotestudo*. The positions of the peripheral plates have been identified but only for *Indotestudo* remains. The peripheral plates for which it was not possible to give a position have been posteriorly assigned to the different ranks following the distribution of those for which a position was determined. Taphonomic alterations have been identified following the atlas of Fernàndez-Jalvo and Andrews (2016). Regarding the size estimations (see below), the mean of the obtained size estimation is considered in the case several measurements were recorded on a single bone. Chi² tests were performed on the Microsoft Excel software 2007 version and other tests with the basic library Stats of the open-source software R (R Core Team, 2020).

### Size estimation of archaeological Indotestudo

In order to reliably estimate the body size of the archaeological individuals of *Indotestudo* sp., we built size estimation equations based on the models of what was previously done for Southeast Asian monitor lizards (Bochaton et al., 2019a), and recently on the size and weight of species of tortoises (Codron et al., 2022; Esker et al., 2019). These approaches are more powerful than considering isolated measurements (e.g., Klein and Cruz-Uribe 1983) to describe the size of subfossil animal populations because: 1) they enable us to take into account several measurements from different anatomical parts to reconstruct the body size structure of a past population, and 2) they convert measurements taken on the skeleton into a variable used to describe the size of modern individuals, which makes it easier to make comparisons needed to address biological questions. To build the equations, we defined a set of 86 measurements (See Supplementary Table 1; Appendix 1) that we recorded on a sample of 34 museum specimens of *Indotestudo* sp. from the Florida Museum of Natural History (UF) and the Comparative Anatomy collection of the Muséum national d’Histoire naturelle (MNHN-ZA-AC) (see all details in Supplementary Table 1). In order to have enough specimens to produce relevant and reliable predictive equations, we pooled together the three species currently included in the genus *Indotestudo*: *I. forstenii* (Schlegel & Müller, 1845): 8 specimens, *I. travancorica* (Boulenger, 1907): 14 specimens, and *I. elongata* (Blyth, 1854): 12 specimens. Differences of body proportions among species were controlled before applying this strategy to avoid including bias related to interspecific differences in the size estimations.

The measurements recorded were taken on all the bones of the plastron and the carapace as well as on the long bones. Vertebrae and skull elements were not taken into account as their occurrences were too rare in the archaeological record. In addition to these measurements, in order to be able to choose a variable accounting for the “body size” of each individual, we took three measurements considered as “size variables” on the complete carapace of the modern specimens: the Carapace Straight Length (CSL), the Shell Height (SH), and the Plastron Length (PL). All the measurements collected on modern and archaeological specimens were recorded using a digital dial caliper [IP 67 (Mitutoyo Corporation, Japan)]. All the measurements recorded are included in the Supplementary Table 1. All statistical analyses were performed using the basic library Stats of the open-source software R (R Core Team, 2020). Each size estimation equation produced is the result of a linear regression between a given log-transformed measurement recorded on a bone/plate and the log-transformed “size variable” of the specimens. The variables are log-transformed to make linear the simple allometric relationship between the used variables (Gould, 1966; Huxley, 1932). Consequently, the obtained CSL estimation has to be log-reversed using an exponential function to be obtained in the same unit as the used measurements. Obtained equations are of the form:

Log (“size variable”) = (Beta1) * log (osteological measurement in mm) + (Beta0)

From this initial set of equations, we chose to discard all the equations that were not significant (p-value above 0.01) and/or with a low coefficient of determination (R2) (below 0.85) in order to keep only the best equations to estimate the size of the archaeological individuals.

### Specific identification of the *I. elongata* archaeological bone sample

The archaeological bones attributed to *I. elongata* have been identified based on a direct comparison with pictures of the skeletal specimens of this species used to build the SCL estimation equations. The family of Testudinidae is only represented by very few species in Southeast Asia (*Geochelone platynota*, *Indotestudo elongata*, *Manouria emys*, and *Manouria impressa*) (Das, 2010) with different sizes, morphologies, and distribution areas, allowing for relatively easy attribution of nearly all of the studied Testudinidae remains to *I. elongata*. Among the *Indotestudo* genus, as the only currently available qualitative diagnostic criterion for *I. elongata* is located on the nuchal plate (presence of a long and narrow cervical scute), we based most of our identification obtained from other plates/bones on the exact similarity between the archaeological bone remains and the morphologies present on the modern specimens of different ages we observed. An overview of the carapace morphologies of juvenile and adult *I. elongata* is provided here (Fig. 1).

**Figure 1.**
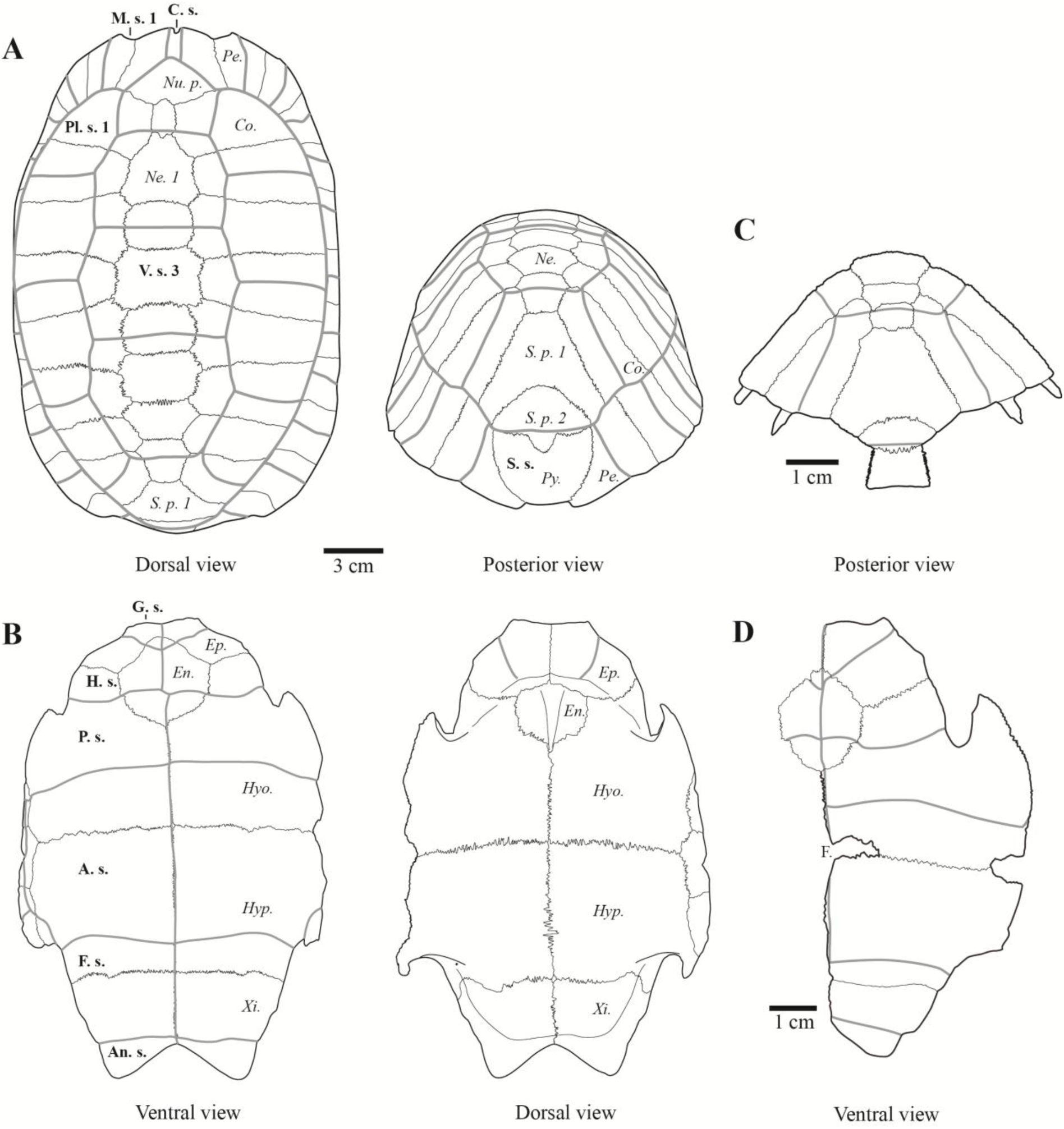
- A) Drawing of the carapace of an adult specimen of *Indotestudo elongata* (CUMZ-R-TT181); B) Drawing of the plastron of an adult specimen of *Indotestudo elongata* (CUMZ-R-TT181); C) Drawing of the carapace of a juvenile specimen of *Indotestudo elongata* lacking peripherals (UF-34760); D) Drawing of the plastron of a juvenile specimen of *Indotestudo elongata* showing central fontanel (UF-34760). **Abbreviations:** A. s.: Abdominal scute, An. s.: Anal scute, G. s.: Gular scute, Co.: Costal plate, F. s.: Femoral scute, Ep.: Epiplastron, En.: Entoplastron, F.: plastral fontanel, H. s.: Humeral scute, Hyo.: Hyoplastron, Hyp.: Hypoplastron, M. s.: Marginal scute, Ne.: Neural plate, Nu. p.: Nuchal plate, C. s.: Cervical scute, P. s.: Pectoral scute, Pe.: Peripheral plate, Pl. s.: Pleural scute, Py.: Pygal plate, S.p. 1: Supra-pygal 1, S.p. 2: Supra-pygal 2, S. s.: Supra-caudal scute, V. s.: Vertebral scute, Xi.: Xiphiplastron.

The remains attributed to this species also present the morphological traits common to most Testudinidae: a carapace lacking lateral keels, a costal pattern of odd costal plates with short distal ends and long medial ends, and even costal plates with long distal ends and short medial ends, octagonal and squared neural plates, peripheral plates without musk ducts, a costo-marginal sulcus superimposed to the costo-peripheral suture, a pygal plate not intersected by the posterior sulcus of the fifth vertebral scute, thickened epiplastra, and thin and vertical inguinal and axillary buttresses. Among Testudinidae, the genus *Indotestudo* is characterized by the fact that the humeropectoral sulcus is crossing the entoplastron (Auffenberg, 1974).

The establishment of robust and quantified diagnostic criteria for the identification of isolated bones of Southeast Asian turtles has yet to be performed. As a comment, we note that the characteristic cervical scute morphology of *I. elongata* was present on all the nuchal plates (N=109) attributed to this species, with the exception of a single remain from Laang Spean cave. It has been previously noted that the cervical scute could be absent on some specimens (Ihlow et al., 2016), and our data indicate a frequency of such a feature less than 1%.

The other families of turtles (Geoemydidae and Trionychidae) have been identified following the diagnostic criteria of Pritchard et al. (2009).

## Results

### Size predictive equations

To determine the most appropriate “size” variable, we conducted correlation tests among the three “body size” measurements recorded on the complete carapaces of our modern specimens of *Indotestudo* spp. The results revealed a strong correlation between the Straight Carapace Length (SCL) and the Plastron Length (PL) (R²=0.97), while the correlation between Shell Height and the other two measurements was weaker (R²=0.93 and 0.92). These differences in correlations could be related to sex-specific or interspecific variations in carapace height among the considered specimens. However, due to the limited size of our sample, we were unable to test these hypotheses conclusively. Consequently, we selected the SCL as our size scalar, although the PL could have been equally considered.

From our complete modern sample, we generated a set of 86 equations, each corresponding to one of the initially recorded measurements. To refine this set and retain the most reliable and precise SCL estimations, we selected 52 equations with significant linear relationships (p-value <0.01) and high coefficients of determination (R²) (Fig. 2; Table. 1). These equations encompass measurements from various anatomical elements, including the epiplastron, entoplastron, hyoplastron, hypoplastron, xiphiplastron, nuchal plate, neural plates (ranks 1, 2, 3, 4, 6, and 7), peripheral plates (ranks 1, 2, 3, 8, 9, and 10), 2nd supra-pygal plate, pygal plate, humerus, radius, ulna, femur, tibia, and fibula.

**Figure 2.** - Measurements corresponding to the 52 equations retained to predict the SCL of our archaeological sample of *Indotestudo elongata* bone remains. Measurements names: GdapW: Greatest distal antero-posterior Width, GdlmW: GH: Greatest Height (on the dorso-ventral axis), GL: Greatest Length (on the antero-posterior axis), GmL: Greatest medial Length (on the antero-posterior axis), GpapW: Greatest proximal antero-posterior Width, GpdvW: Greatest proximal dorso-ventral Width, GplmW: Greatest proximal latero-medial Width, GW: Greatest Width (on the latero-medial axis).

### Zooarchaeological and taphonomic analyses of the herpetofaunal assemblages

#### Doi Pha Kan Rockshelter

##### Composition of the herpetofaunal assemblages

The herpetofaunal assemblage of Doi Pha Kan consists of 8414 bone remains, weighing a total of 6875 grams, and representing at least 115 individuals. The majority of these bones belong to non-marine turtles (Fig. 3; Table. 2), accounting for 74% of the Weight of the Remains (WR), 56% of the Number of Identified Specimens (NISP), and 47% of the Minimum Number of Individuals (MNI). The second most represented group is Monitor lizards comprising 23% of the WR, 38% of the NISP, and 38% of the MNI. There are also bone remains from rare snakes, amphibians, and small lizards, although they are less common in the assemblage.

**Figure 3.**
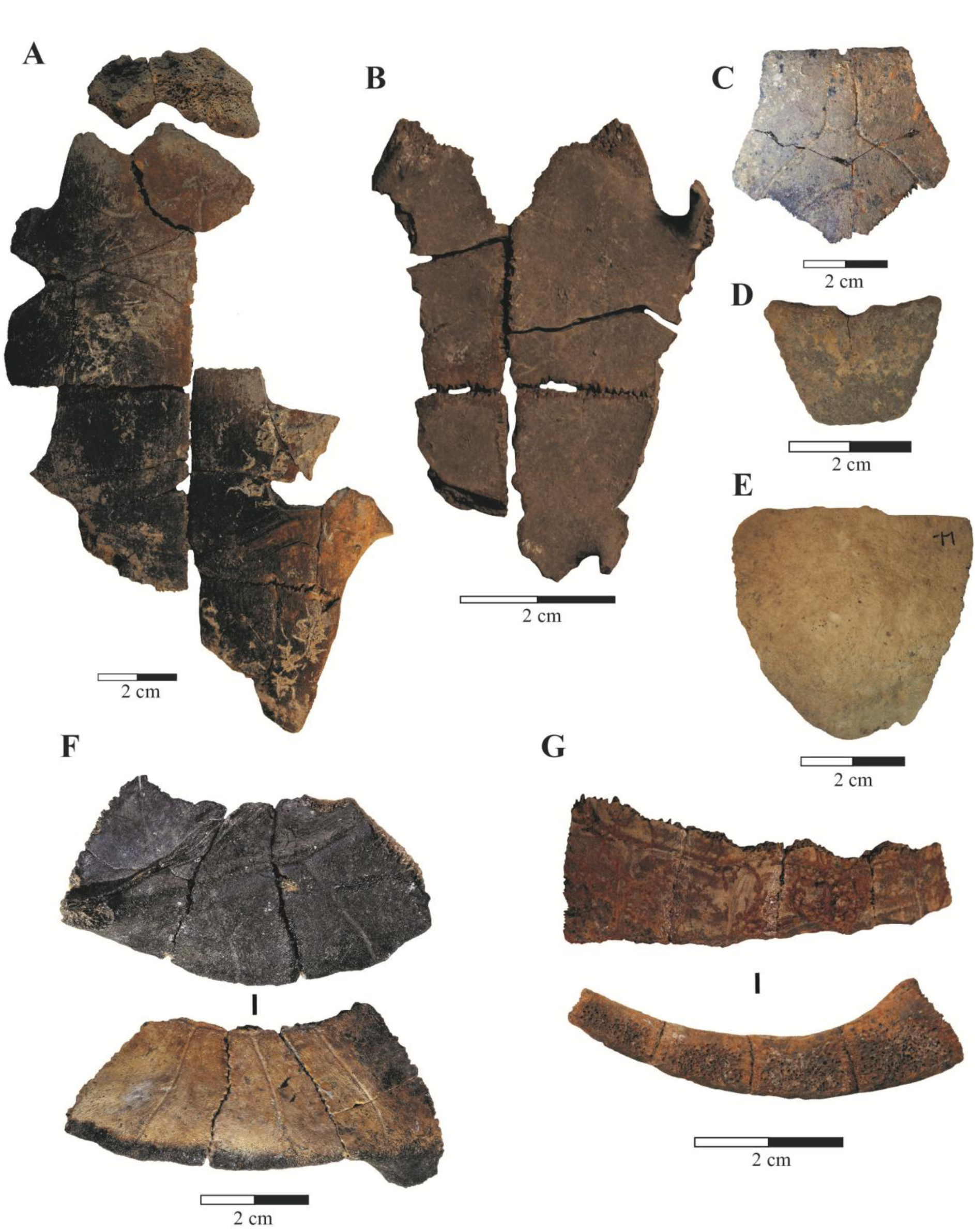
- Examples of the studied turtle bone remains: A) Plastron of an adult *I. elongata* in anatomical connection from the Hoabinhian layer of Laang Spean Cave (ventral view); B) Plastron of a juvenile *I. elongata* in anatomical connection from the Hoabinhian layer of Laang Spean Cave (ventral view); C) Nuchal plate of *I. elongata* from the Hoabinhian layer of Laang Spean Cave (dorsal view); D) Pygal plate of a juvenile *I. elongata* from the Neolithic layer of Laang Spean Cave (posterior view); E) Pygal plate of an adult *I. elongata* from the Neolithic layer of Laang Spean Cave (posterior view); F) Left peripheral plates (1^st^ to 3^rd^) of *I. elongata* in anatomical connection from layer 2 of Moh Khiew Cave showing burning traces limited to the internal side of the carapace; G) Left peripheral plates (8^th^ to 11^th^) of a Geoemydidae in anatomical connection with the ventral part cut-down from the site of Doi Pha Kan.

In terms of the bones attributed to turtles and tortoises (Table. 3), a significant portion, 66.8% (50.8% of the WR), could not be identified to a specific family level. Among the fragments that were identified to at least the family level, the majority (71%) were attributed to *Indotestudo elongata* (71% of the WR and 76% of the MNI), while 28% (28% of the WR and 18% of the MNI) were assigned to the family Geoemydidae. Only a small percentage (less than 1%) of the remains, were attributed to the family Trionychidae (0.6% of the WR and 5% of the MNI). It is worth noting that Trionychid remains can be identified for all the plates due to their ornamentation, so they are not underrepresented in the assemblage simply because they are difficult to identify.

##### Taphonomy of the turtle/tortoise bone assemblage

Among the 4762 bone fragments attributed to turtles/tortoises, 303 are complete elements (6.3%), and 249 are nearly complete (at least 90% of the bone is preserved), while 1389 (29%) are small fragments representing less than 5% of the complete anatomical part. The average percentage of completion of the bones is 32%. The overall Percentage of Representation (PR) is 28%. The best-represented bones (Fig. 4 – A) are the stylopods (humerus and femur) with PR>75%, followed by the tibia, radius, scapula, coracoid, and epiplastron with PR>50%. The rest of the long bones and most of the easily identifiable plates also have relatively high representation. However, the peripheral plates of the bridge are less represented (PR=10%) compared to all the other peripheral plates (PR=39%). The skull, vertebrae, and all small elements of the hands and feet are nearly absent. The largest plates (hyoplastron and hypoplastron) are the most fragmented, with completion means of less than 26%. The peripheral plates corresponding to the bridge are also more heavily fragmented (completion mean of 59%) compared to the other peripheral plates (completion means 67-89%). Traces of burning (black) and carbonization (grey/white) were observed on 690 bones (14% of the total NISP). These traces were present indiscriminately on every anatomical element of the carapace and skeleton. Cut marks were observed on only five bones, six peripheral plates, and one hypoplastron. Notably, a series of peripheral plates attributed to a Geoemydidae turtle bears clear traces of a clean cut aiming to cut the ventral part of the carapace (Fig. 3-G). Lastly, 120 bones have been observed as corresponding to preserved anatomical connections on the field between unfused plates. This indicates that at least a part of the assemblage was undisturbed prior to the excavation.

**Figure 4.**
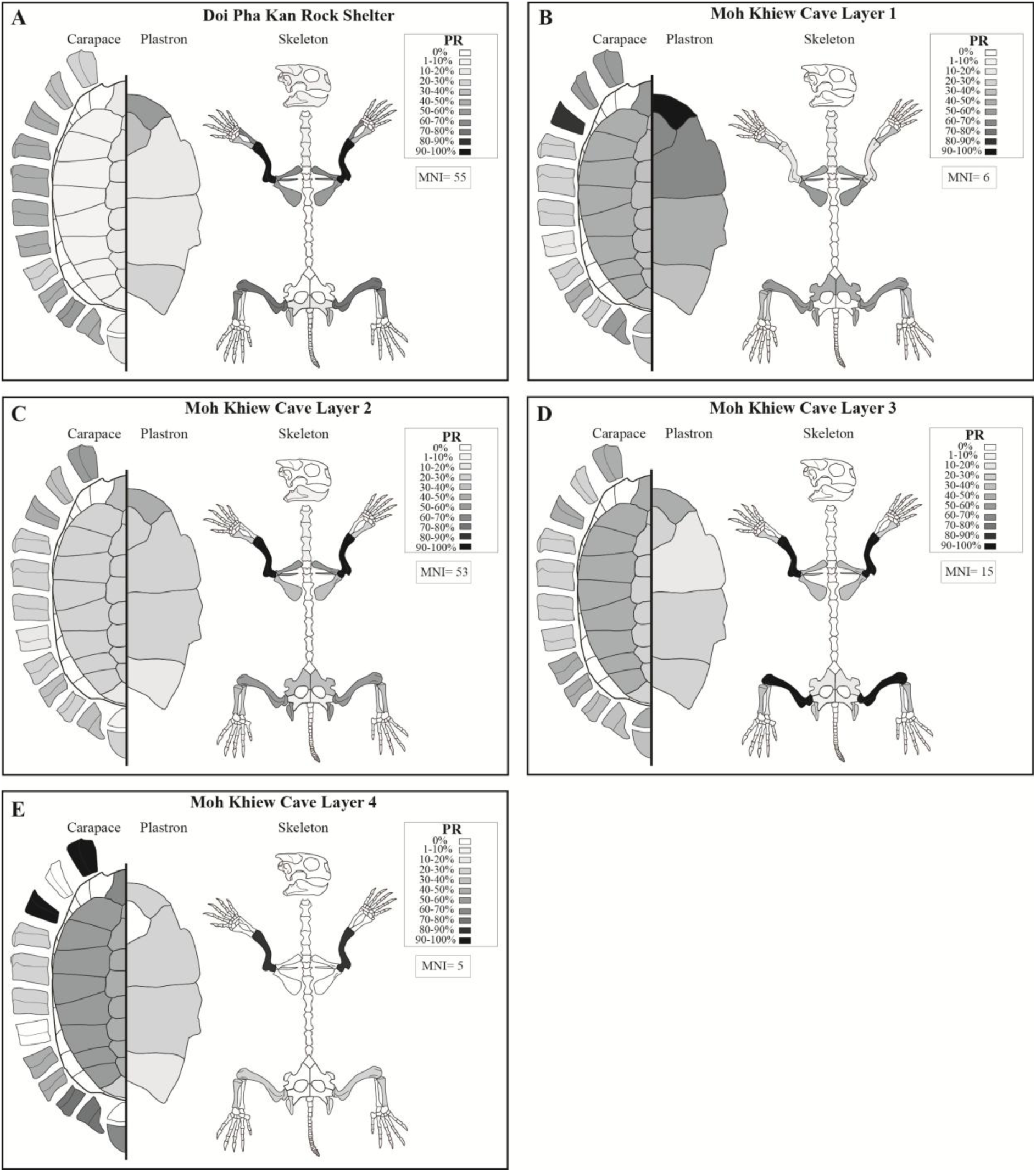
- Anatomical distributions of the turtle/tortoise remains collected in the sites of Doi Pha Kan rockshelter (A), and Moh Khiew Cave (B-E). The percentage of representation (PR) is considered here to provide a graphical visualization of the different values observed for the different anatomical elements.

##### Size of *Indotestudo elongata* archaeological individuals

The measurements recorded on the archaeological material of Indotestudo elongata from Doi Pha Kan allowed for the reconstruction of 201 Straight Carapace Length (SCL) estimations, ranging from 64 to 292 mm, with a mean SCL of 182 mm (Figure. 5-A). These estimations correspond to at least 42 individuals. The distribution of these sizes was found to be non-unimodal based on Hartigans’ dip test (p. value > 0.05), suggesting the possibility of a bimodal distribution with one population of small individuals around 120 mm and a second one ranging from 140 to 260 mm.

**Figure 5.**
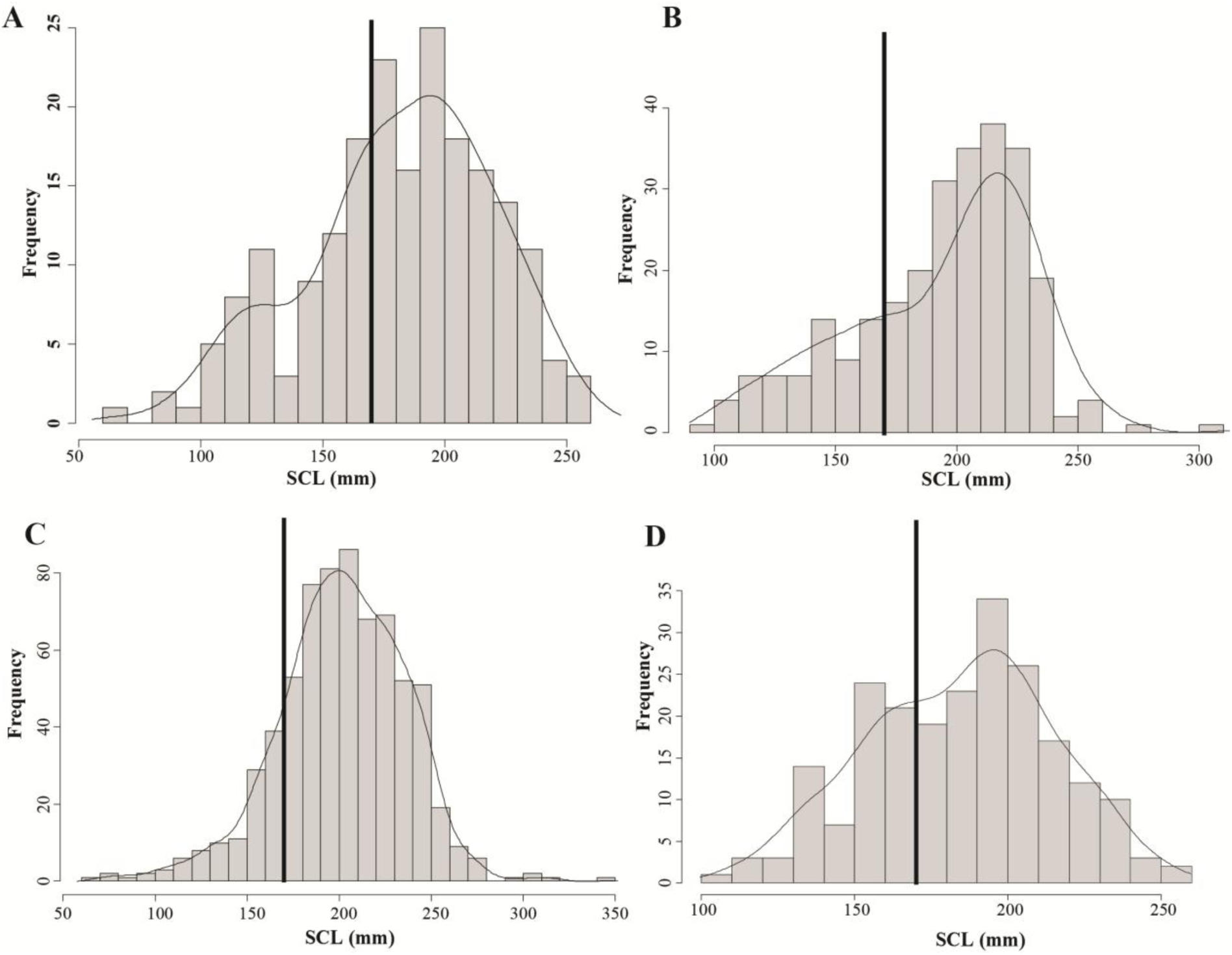
- Histograms of the *Indotestudo elongata* size reconstructions (Standard Carapace Length) obtained from the different herpetofaunal bone assemblages studied: A) Doi Pha Kan Rock-shelter (NMI=42); B) Moh Khiew Cave (NMI=59); C) Laang Spean Cave (NMI=75); D) Khao Ta Phlai Cave (NMI=26). The black bars represent the smallest size of the sexually mature specimens based on modern data collected on modern *I. elongata* populations.

#### Moh Khiew cave

##### Composition of the herpetofaunal assemblages

The herpetofaunal assemblage of Moh Khiew Cave consists of 9,108 bone remains weighing 8,351 grams. These bones are mainly distributed in layers 2 (51% of the NISP and 52% of the WR), 1 (26% of the NISP and 24% of the WR), and 3 (17% of the NISP and 18% of the WR) (Table. 4). The complete assemblage includes bone fragments from at least 152 individuals.

The majority of the sample corresponds to non-marine turtle remains (63% of the NISP, 74% of the WR, and 52% of the MNI), followed by Monitor lizards (25% of the NISP, 16% of the WR, and 24% of the MNI), and snakes (8% of the NISP, 8% of the WR, and 4% of the MNI). The remains of small-sized lizards (excluding snakes), amphibians, and crocodiles altogether represent less than 5% of the assemblage in terms of NISP and WR. However, the distribution of these groups among the layers is not homogeneous. Turtle bone remains represent more than 75% of the NISP, 85% of the WR, and 60% of the MNI in layers 2 to 4, but only 20% of the NISP, 35% of the WR, and 21% of the MNI in layer 1. On the other hand, Monitor lizards represent 46% of the NISP, 31% of the WR, and 35% of the MNI in the layer 1, but less than 20% of NISP, 13% of the WR, and 25% of the MNI in the other layers. Snakes are also better represented in layer 1 (23% of the NISP) than in the subsequent levels (less than 2.6% of the NISP). Chi² tests performed on the NISP indicate that the faunal composition of layer 1 significantly differs (P. value < 0.01) from the layers 2 and 3, while the sample size of layer 4 is too low to conduct a statistical test.

Regarding the identification of non-marine turtle taxa (Table 5), between 66% and 79% of the bone fragments have not been associated with at least a family rank identification. Among the identified families, Testudinidae (*Indotestudo elongata*) represents a consistently major part of the identified bone remains, ranging from 68% to 63% of the NISP, 70% to 56% of the WR, and 86% to 50% of the MNI depending on the layer. The other fragments have been attributed to Geoemydidae turtles, accounting for 36% to 29% of the NISP, 36% to 28% of the WR, and 33% to 12% of the MNI, while Trionychidae comprises only 0% to 2% of the NISP and WR. The composition of the turtle assemblages in terms of families seems to be fairly stable across the layers, as indicated by the results of Chi² tests, which did not show significant differences in the distribution of the NISP from the different testable layers (p. value > 0.01).

**Table 1.**
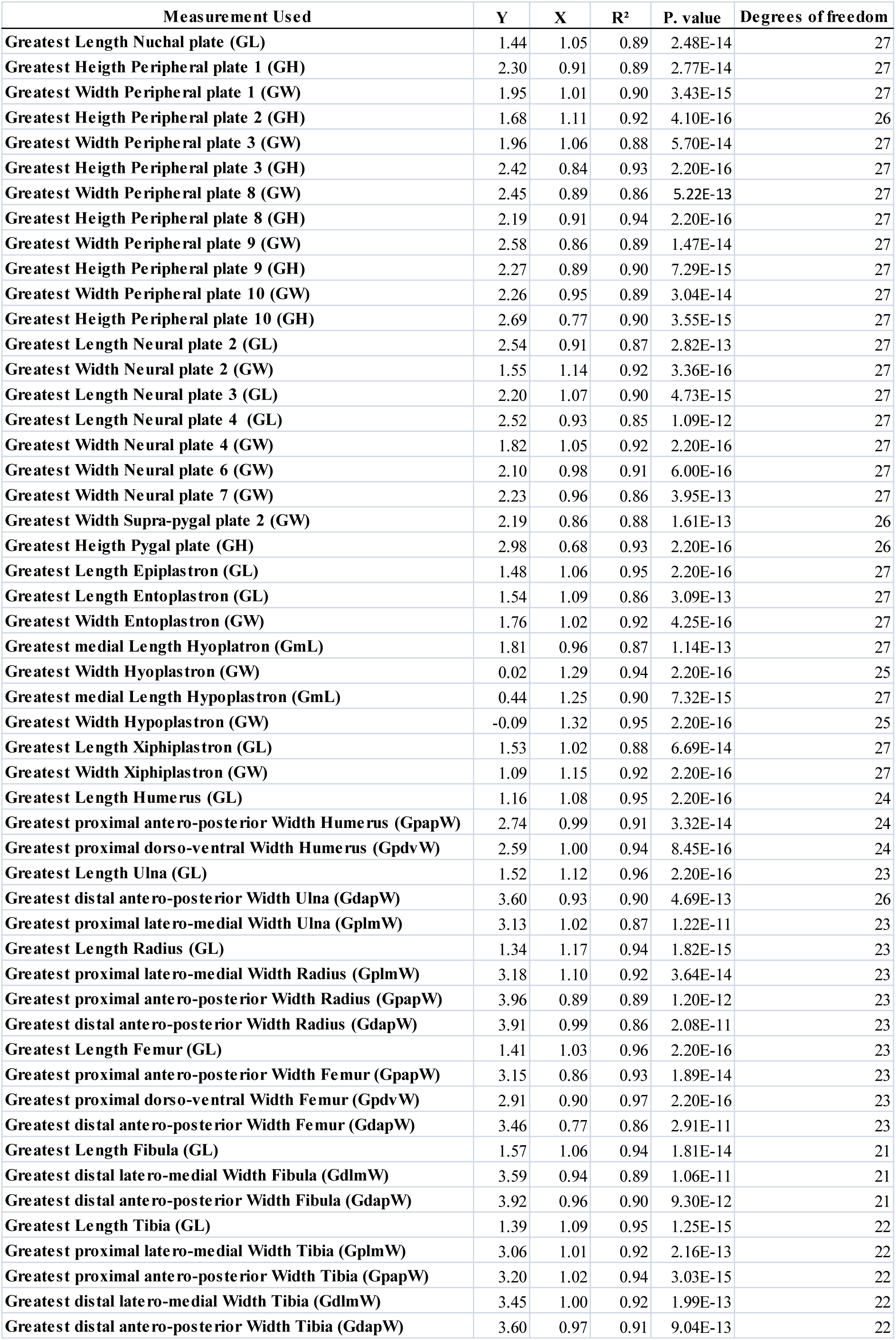
- Equations retained for predicting the Straight Carapace Length (SCL) of the archaeological Indotestudo specimens. Are indicated: the measurement used (to be recorded on the archaeological specimen), the slope X (“Beta1” to integrate into the equation as mentioned in the Material and Methods section), the intercept Y (“Beta0” to integrate into the equation as mentioned in the Material and Methods section), the coefficient of determination (R²) of each relationship, the p-values, and the degrees of freedom for each linear regression.

| Measurement Used | Y | X | R<sup>2</sup> | P. value | Degrees of freedom |
| --- | --- | --- | --- | --- | --- |
| Greatest Length Nuchal plate (GL) | 1.44 | 1.05 | 0.89 | 2.48E-14 | 27 |
| Greatest Height Peripheral plate 1 (GH) | 2.30 | 0.91 | 0.89 | 2.77E-14 | 27 |
| Greatest Width Peripheral plate 1 (GW) | 1.95 | 1.01 | 0.90 | 3.43E-15 | 27 |
| Greatest Height Peripheral plate 2 (GH) | 1.68 | 1.11 | 0.92 | 4.10E-16 | 26 |
| Greatest Width Peripheral plate 3 (GW) | 1.96 | 1.06 | 0.88 | 5.70E-14 | 27 |
| Greatest Height Peripheral plate 3 (GH) | 2.42 | 0.84 | 0.93 | 2.20E-16 | 27 |
| Greatest Width Peripheral plate 8 (GW) | 2.45 | 0.89 | 0.86 | 5.22E-13 | 27 |
| Greatest Height Peripheral plate 8 (GH) | 2.19 | 0.91 | 0.94 | 2.20E-16 | 27 |
| Greatest Width Peripheral plate 9 (GW) | 2.58 | 0.86 | 0.89 | 1.47E-14 | 27 |
| Greatest Height Peripheral plate 9 (GH) | 2.27 | 0.89 | 0.90 | 7.29E-15 | 27 |
| Greatest Width Peripheral plate 10 (GW) | 2.26 | 0.95 | 0.89 | 3.04E-14 | 27 |
| Greatest Height Peripheral plate 10 (GH) | 2.69 | 0.77 | 0.90 | 3.55E-15 | 27 |
| Greatest Length Neural plate 2 (GL) | 2.54 | 0.91 | 0.87 | 2.82E-13 | 27 |
| Greatest Width Neural plate 2 (GW) | 1.55 | 1.14 | 0.92 | 3.36E-16 | 27 |
| Greatest Length Neural plate 3 (GL) | 2.20 | 1.07 | 0.90 | 4.73E-15 | 27 |
| Greatest Length Neural plate 4 (GL) | 2.52 | 0.93 | 0.85 | 1.09E-12 | 27 |
| Greatest Width Neural plate 4 (GW) | 1.82 | 1.05 | 0.92 | 2.20E-16 | 27 |
| Greatest Width Neural plate 6 (GW) | 2.10 | 0.98 | 0.91 | 6.00E-16 | 27 |
| Greatest Width Neural plate 7 (GW) | 2.23 | 0.96 | 0.86 | 3.95E-13 | 27 |
| Greatest Width Supra-pygial plate 2 (GW) | 2.19 | 0.86 | 0.88 | 1.61E-13 | 26 |
| Greatest Height Pygal plate (GH) | 2.98 | 0.68 | 0.93 | 2.20E-16 | 26 |
| Greatest Length Epiplastron (GL) | 1.48 | 1.06 | 0.95 | 2.20E-16 | 27 |
| Greatest Length Entoplastron (GL) | 1.54 | 1.09 | 0.86 | 3.09E-13 | 27 |
| Greatest Width Entoplastron (GW) | 1.76 | 1.02 | 0.92 | 4.25E-16 | 27 |
| Greatest medial Length Hyoplastron (GmL) | 1.81 | 0.96 | 0.87 | 1.14E-13 | 27 |
| Greatest Width Hyoplastron (GW) | 0.02 | 1.29 | 0.94 | 2.20E-16 | 25 |
| Greatest medial Length Hypoplastron (GmL) | 0.44 | 1.25 | 0.90 | 7.32E-15 | 27 |
| Greatest Width Hypoplastron (GW) | -0.09 | 1.32 | 0.95 | 2.20E-16 | 25 |
| Greatest Length Xiphiplastron (GL) | 1.53 | 1.02 | 0.88 | 6.69E-14 | 27 |
| Greatest Width Xiphiplastron (GW) | 1.09 | 1.15 | 0.92 | 2.20E-16 | 27 |
| Greatest Length Humerus (GL) | 1.16 | 1.08 | 0.95 | 2.20E-16 | 24 |
| Greatest proximal antero-posterior Width Humerus (GpapW) | 2.74 | 0.99 | 0.91 | 3.32E-14 | 24 |
| Greatest proximal dorso-ventral Width Humerus (GpdvW) | 2.59 | 1.00 | 0.94 | 8.45E-16 | 24 |
| Greatest Length Ulna (GL) | 1.52 | 1.12 | 0.96 | 2.20E-16 | 23 |
| Greatest distal antero-posterior Width Ulna (GdapW) | 3.60 | 0.93 | 0.90 | 4.69E-13 | 26 |
| Greatest proximal latero-medial Width Ulna (GplmW) | 3.13 | 1.02 | 0.87 | 1.22E-11 | 23 |
| Greatest Length Radius (GL) | 1.34 | 1.17 | 0.94 | 1.82E-15 | 23 |
| Greatest proximal latero-medial Width Radius (GplmW) | 3.18 | 1.10 | 0.92 | 3.64E-14 | 23 |
| Greatest proximal antero-posterior Width Radius (GpapW) | 3.96 | 0.89 | 0.89 | 1.20E-12 | 23 |
| Greatest distal antero-posterior Width Radius (GdapW) | 3.91 | 0.99 | 0.86 | 2.08E-11 | 23 |
| Greatest Length Femur (GL) | 1.41 | 1.03 | 0.96 | 2.20E-16 | 23 |
| Greatest proximal antero-posterior Width Femur (GpapW) | 3.15 | 0.86 | 0.93 | 1.89E-14 | 23 |
| Greatest proximal dorso-ventral Width Femur (GpdvW) | 2.91 | 0.90 | 0.97 | 2.20E-16 | 23 |
| Greatest distal antero-posterior Width Femur (GdapW) | 3.46 | 0.77 | 0.86 | 2.91E-11 | 23 |
| Greatest Length Fibula (GL) | 1.57 | 1.06 | 0.94 | 1.81E-14 | 21 |
| Greatest distal latero-medial Width Fibula (GdlmW) | 3.59 | 0.94 | 0.89 | 1.06E-11 | 21 |
| Greatest distal antero-posterior Width Fibula (GdapW) | 3.92 | 0.96 | 0.90 | 9.30E-12 | 21 |
| Greatest Length Tibia (GL) | 1.39 | 1.09 | 0.95 | 1.25E-15 | 22 |
| Greatest proximal latero-medial Width Tibia (GplmW) | 3.06 | 1.01 | 0.92 | 2.16E-13 | 22 |
| Greatest proximal antero-posterior Width Tibia (GpapW) | 3.20 | 1.02 | 0.94 | 3.03E-15 | 22 |
| Greatest distal latero-medial Width Tibia (GdlmW) | 3.45 | 1.00 | 0.92 | 1.99E-13 | 22 |
| Greatest distal antero-posterior Width Tibia (GdapW) | 3.60 | 0.97 | 0.91 | 9.04E-13 | 22 |

**Table 2.** - Weight of the Remains (WR), Number of Identified Specimens (NISP), and Minimum Number of Individuals (MNI) corresponding to the different taxa identified in the complete Doi Pha Kan Rock-shelter herpetofaunal assemblage.

|  | **NISP** | **WR** | **MNI** |
| --- | --- | --- | --- |
| Turtle/Tortoise | 4762 | 5113.02 | 55 |
| Monitor lizard | 3203 | 1610.39 | 44 |
| Snake | 375 | 141 | 3 |
| Amphibian | 67 | 10.57 | 10 |
| Small lizard | 7 | 0.82 | 3 |
| **Total** | **8414** | **6875.8** | **115** |

**Table 3.** - Number of Identified Specimens (NISP), Weight of the Remains (WR), and Minimum Number of Individuals (MNI) corresponding to the different turtle/tortoise taxa identified in the complete Doi Pha Kan Rock-shelter assemblage.

|  | **NISP** | **WR** | **MNI** |
| --- | --- | --- | --- |
| *Indotestudo elongata* | 1122 | 1787.17 | 42 |
| Geoemydidae | 447 | 711.34 | 10 |
| Trionychidae | 12 | 14.13 | 3 |
| Turtle ind. | 3181 | 2600.38 |  |
| **Total** | **4762** | **5113.02** | **55** |

**Table 4.** - Number of Identified Specimens (NISP), Weight of the remains (WR), and Minimum Number of Individuals (MNI) identified in the complete herpetofaunal assemblage of the different layers of the 2008 excavation of Moh Khiew Cave.

|  | Layer 1 |  |  | Layer 2 |  |  | Layer 3 |  |  | Layer 4 |  |  | Total |  |  |
| --- | --- | --- | --- | --- | --- | --- | --- | --- | --- | --- | --- | --- | --- | --- | --- |
|  | NISP | WR | MNI | NISP | WR | MNI | NISP | WR | MNI | NISP | WR | MNI | NISP | WR | MNI |
| Turtle/Tortoise | 503 | 702.8 | 6 | 3614 | 3789 | 53 | 1184 | 1288 | 15 | 439 | 481.3 | 5 | **5740** | **6261** | **79** |
| Monitor lizard | 1111 | 641.4 | 17 | 872 | 463.8 | 12 | 298 | 190.1 | 5 | 43 | 25.4 | 2 | **2324** | **1321** | **36** |
| Snake | 560 | 612.2 | 3 | 117 | 52.8 | 1 | 37 | 29.15 | 1 | 7 | 6.4 | 1 | **721** | **700.6** | **6** |
| Amphibian | 144 | 38.45 | 14 | 34 | 6.5 | 4 | 22 | 3.92 | 3 | 0 | 0 | 0 | **200** | **48.87** | **21** |
| Small lizard | 89 | 8.27 | 7 | 4 | 0.3 | 1 | 28 | 3.1 | 1 | **0** | **0** | **0** | **121** | **11.67** | **9** |
| Crocodile | 2 | 8.5 | 1 | 0 | 0 | 0 | 0 | 0 | 0 | 0 | 0 | 0 | **2** | **8.5** | **1** |
| **Total** | **2409** | **2012** | **48** | **4641** | **4312** | **71** | **1569** | **1514** | **25** | **489** | **513.1** | **8** | **9108** | **8351** | **152** |

**Table 5.**
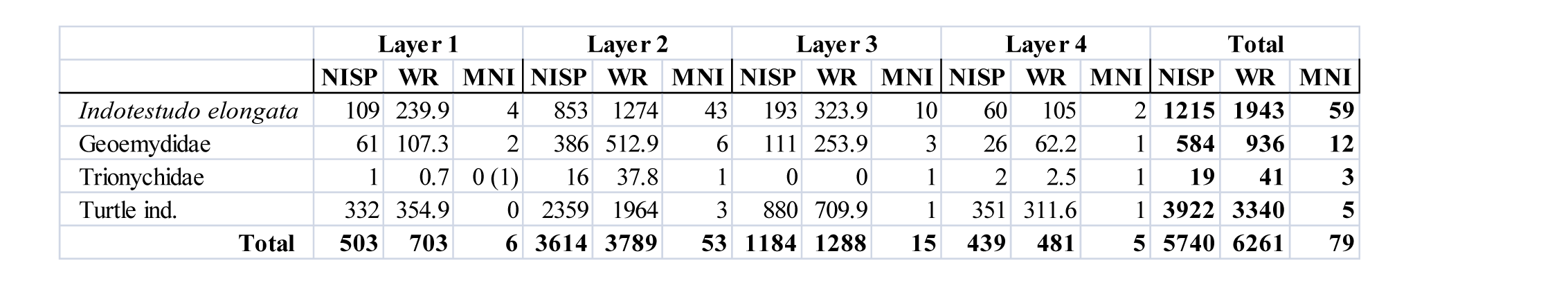
- Number of Identified Specimens (NISP), Weight of the remains (WR), and Minimum Number of Individuals (MNI) in the non-marine turtle assemblage from the different layers of the 2008 excavation of Moh Khiew Cave.

|  | Layer 1 |  |  | Layer 2 |  |  | Layer 3 |  |  | Layer 4 |  |  | Total |  |  |
| --- | --- | --- | --- | --- | --- | --- | --- | --- | --- | --- | --- | --- | --- | --- | --- |
|  | NISP | WR | MNI | NISP | WR | MNI | NISP | WR | MNI | NISP | WR | MNI | NISP | WR | MNI |
| *Indotestudo elongata* | 109 | 239.9 | 4 | 853 | 1274 | 43 | 193 | 323.9 | 10 | 60 | 105 | 2 | **1215** | **1943** | **59** |
| Geoemydidae | 61 | 107.3 | 2 | 386 | 512.9 | 6 | 111 | 253.9 | 3 | 26 | 62.2 | 1 | **584** | **936** | **12** |
| Trionychidae | 1 | 0.7 | 0 (1) | 16 | 37.8 | 1 | 0 | 0 | 1 | 2 | 2.5 | 1 | **19** | **41** | **3** |
| Turtle ind. | 332 | 354.9 | 0 | 2359 | 1964 | 3 | 880 | 709.9 | 1 | 351 | 311.6 | 1 | **3922** | **3340** | **5** |
| **Total** | **503** | **703** | **6** | **3614** | **3789** | **53** | **1184** | **1288** | **15** | **439** | **481** | **5** | **5740** | **6261** | **79** |

**Table 6.** - Number of Identified Specimens (NISP), Weight of the Remains (WR), Number of Remains (NR), and Minimum Number of Individuals (MNI) corresponding to the different herpetofaunal taxa identified in the herpetofaunal bone assemblages collected in the two test-pits of the site of Khao Ta Phlai.

|  | TP1 |  |  | TP2 |  |  |
| --- | --- | --- | --- | --- | --- | --- |
|  | NISP | WR | MNI | NISP | WR | MNI |
| Amphibian |  |  |  | 3 | 0.51 | 1 |
| Snake | 38 | 33.4 | 2 | 37 | 69.3 | 2 |
| Turtle/Tortoise | 1335 | 2509.7 | 19 | 1825 | 3901 | 18 |
| Monitor lizard | 274 | 347.7 | 6 | 251 | 377.9 | 6 |
| **Total** | **1647** | **2891** | **27** | **2116** | **4349** | **27** |

##### Taphonomy of the turtle/tortoise bone assemblage

The fragmentation of the 5740 bone fragments analyzed increases with depth. In layer 1, 8.9% of the bones are complete, and 12.9% are nearly complete. In layer 2, these percentages fall to 7.5% and 12.1%, then to 5.6% and 9.6% in layer 3, and finally to 3.9% and 7.7% in layer 4. The average percentage of completeness of the bones is also slightly higher in layers 1 and 2 (36% and 37%) than in layers 3 and 4 (34% and 28%). These differences are only significant between layers 1 and 4 (Chi² test; p. value < 0.01).

The anatomical distribution of the remains shows strong variations between the layers (Fig. 4), but the sizes of the assemblages are also very dissimilar, with the bone samples of layers 1 and 4 containing only 503 and 439 remains, respectively, while those of layers 2 and 3 contain 3614 and 1184 bone fragments. Distributions in small samples could be more strongly impacted by random effects than larger assemblages, and a strict comparison of the four layers might not make sense at all. Some general trends can, however, still be noted, such as the overall PR, which is between 27% and 24% in all the layers, except the first in which it is slightly higher (36%). Nearly all anatomical parts are present in every layer, but the skulls, vertebrae, and extremities are nearly absent. The stylopods (humerus and femur) are the best-represented bones in the richest layers. They are also well-represented in layers 1 and 2 but are outnumbered by some specific carapace plates. Girdles and zeugopods are also present but in smaller numbers. Regarding the carapace and the plastron, no clear pattern emerges except for the nearly systematic lower representation of the peripheral plates of the bridge (PR=25%-21%) compared to the other peripheral plates (mean PR=52%-29%). This could be attributed to an identification bias stemming from the comparatively lower mean completion rate of peripheral plates of the bridge, potentially making their identification more challenging compared to other peripherals.

Burning (black) and carbonization (grey/white) traces were observed on 325 bones (5.7% of the total NISP). Such traces were present indiscriminately on every anatomical element and every side of the carapace and skeleton parts. They were recorded on the internal side of several peripheral plates that were still in anatomical connection at the moment of the excavation (Fig. 3-F). However, such observations were not repeated on the rest of the material. Cut marks were observed on only three bones: one peripheral plate, one nuchal plate, and one xiphiplastron. Among the full assemblage, 61 fragments of carapaces were still in anatomical connection at the moment of the excavation. These elements were distributed mostly in layers 2 and 3 but also in the lower part of layer 1 at a depth of 70-80 cm.

##### Size of *Indotestudo elongata* individuals

The measurements recorded on the *I. elongata* archaeological material of Moh Khiew Cave enabled the reconstruction of 201 SCL estimations, ranging from 98 to 310 mm, with a mean of 193 mm (Figure. 5-B), corresponding to at least 59 individuals. The mean size of the tortoises is similar in all layers, and no statistically significant differences were noted (student T-test; p. value > 0.01). In layer 1 (N=25), the mean size was 198 mm, 194 mm in layer 2 (N=169), 187 mm in layer 3 (N=52), and finally 195 mm in layer 4 (N=19). The largest observed specimen was in layer 1. The global distribution of these sizes was not unimodal (Hartigans’ dip test, p. value > 0.01), and mixture models indicate it is most likely bimodal, with a best-represented group of individuals around 220 mm and a second group of smaller specimens around 150 mm.

#### Khao Ta Phlai Cave

##### Taxonomic composition

A total of 3763 bone remains of herpetofauna, weighing 7239 grams and representing at least 43 individuals, were analyzed from the two excavated test-pits of the site of Khao Ta Phlai (Table. 6). Most of these bones correspond to turtles or tortoises in terms of WR (87% in TP1 and 90% in TP2), NISP (81% in TP1 and 86% in TP2), and MNI (65% in TP1 and 70% in TP2). The second most represented herpetofaunal group is the Monitor lizards (12% and 9% of the WR, 16% and 12% of the NISP, and 23% and 22% of the MNI) followed by some rare remains of snakes, as well as a few amphibian bones in TP2 only.

Although the distribution of the material between the two TPs is somewhat homogeneous, the repartition of the bones across the two main periods documented (Metal Ages and Neolithic) is quite different. In TP2, most of the material (95% of the NISP) is located in the Neolithic layers, while in TP1, the bones are more evenly distributed (55% of the NISP in the Metal Ages layers and 44% of the NISP in the Neolithic levels).

The distribution of the main herpetofaunal taxa across the layers does not present strong variations (Tab. 7), as only the very poor Metal Ages layer of TP2 significantly differs from the other levels (Chi²; p. value < 0.01). Turtle bones are always the best represented (between 71% and 87% of the NISP), but Monitor lizards seem to be slightly better represented in Metal Ages layers from TP1 and 2 (18% and 20% compared to Neolithic layers (14% and 11% of the NISP). These tendencies are, however, not statistically significant and difficult to interpret in the absence of a study of the complete faunal assemblages.

**Table 7.** - Number of Identified Specimens (NISP), Weight of the Remains (WR), and Minimum Number of Individuals (MNI) corresponding to the different herpetofaunal taxa identified in the herpetofaunal bone assemblages of the two periods identified in the two test-pits of the site of Khao Ta Phlai.

|  | TP1-Metal Age |  |  | TP1-Neolithic |  |  | TP2-Metal Age |  |  | TP2-Neolithic |  |  |
| --- | --- | --- | --- | --- | --- | --- | --- | --- | --- | --- | --- | --- |
|  | NISP | WR | MNI | NISP | WR | MNI | NISP | WR | MNI | NISP | WR | MNI |
| Amphibian |  |  |  |  |  |  |  |  |  | 3 | 0.51 | 1 |
| Snake | 33 | 22.4 | 2 | 5 | 11 | 2 | 9 | 13.4 | 2 | 28 | 55.9 | 2 |
| Turtle/Tortoise | 720 | 1011.8 | 14 | 615 | 1497.8 | 9 | 70 | 131.2 | 2 | 1755 | 3769.8 | 21 |
| Monitor lizard | 166 | 146.2 | 1 | 108 | 201.5 | 1 | 20 | 36.2 | 1 | 231 | 341.7 | 3 |
| **Total** | **919** | **1180** | **17** | **728** | **1710** | **12** | **99** | **180.8** | **5** | **2017** | **4168** | **27** |

The identification rate of non-marine turtle bones was lower in TP1 (41% of the WR and 25% of the NR) than in TP2 (57% of the WR and 35% of the NR). Regarding the bones attributed to a given family (Tab. 8), TP1 provided nearly as much Testudinidae as Geoemydidae in terms of WR and NISP, while *Indotestudo elongata* is much more represented than the latter in TP2 (61% of the WR and 69% of the NISP). These differences are statistically significant (Chi² test; p. value < 0.01). Trionychidae are present in the two TPs.

**Table 8.** - Number of Identified Specimens (NISP), Weight of the Remains (WR), and Minimum Number of Individuals (MNI) corresponding to the different turtle/tortoise taxa identified in the bone assemblages collected in the two test-pits of the site of Khao Ta Phlai.

|  | TP1 |  |  | TP2 |  |  |
| --- | --- | --- | --- | --- | --- | --- |
|  | NISP | WR | MNI | NISP | WR | MNI |
| *Indotestudo elongata* | 175 | 435.5 | 8 | 441 | 1357 | 14 |
| Geoemydidae | 150 | 581.9 | 5 | 191 | 838.6 | 3 |
| Trionychidae | 10 | 14.1 | 1 | 9 | 32.2 | 1 |
| Turtle ind. | 1000 | 1478 |  | 1184 | 1673 |  |
| **Total** | **1335** | **2510** | **14** | **1825** | **3901** | **18** |

If the chronological phases are considered (Tab. 9), Geoemydidae and Trionychidae are significantly better represented in the upper layers of TP2 and TP1, corresponding to the Metal Ages. These two layers do not significantly differ in terms of family composition (Chi² test; p. value > 0.01) but significantly differ from the two Neolithic layers (Chi² test; p. value < 0.01). This trend of a more important exploitation of freshwater turtles during the Metal Ages in regard to the Neolithic period is, for now, difficult to interpret considering the possible issues of chronological associations between the layers of the two TPs and the possibility of spatial variation in the distribution of the remains inside the site. Indeed, freshwater turtles are also better represented in the Neolithic layer of TP1 compared to TP2.

**Table 9.** - Number of Identified Specimens (NISP), Weight of the Remains (WR), and Minimum Number of Individuals (MNI) corresponding to the different turtle/tortoise taxa identified in the bone assemblages collected in the different chronological phases of the two test-pits of the site of Khao Ta Phlai.

|  | TP1-MA |  |  | TP1-NE |  |  | TP2-MA |  |  | TP2-NE |  |  |
| --- | --- | --- | --- | --- | --- | --- | --- | --- | --- | --- | --- | --- |
|  | NISP | WR | MNI | NISP | WR | MNI | NISP | WR | MNI | NISP | WR | MNI |
| *Indotestudo elongata* | 86 | 149 | 7 | 89 | 286.5 | 6 | 14 | 40.2 | 1 | 427 | 1316.6 | 12 |
| Geoemydidae | 81 | 243.2 | 2 | 69 | 338.7 | 3 | 13 | 28.7 | 1 | 178 | 809.9 | 4 |
| Trionychidae | 10 | 14.1 | 1 |  |  |  | 3 | 5.2 | 1 | 6 | 27 | 1 |
| Turtle ind. | 543 | 605 |  | 457 | 872.63 |  | 70 | 57.1 |  | 1144 | 1616.3 |  |
| **Total** | **720** | **1011** | **10** | **615** | **1498** | **9** | **100** | **131.2** | **3** | **1755** | **3770** | **17** |

##### Taphonomy of the turtle/tortoise bone assemblage

Among the 3, 160 bone fragments attributed to turtle/tortoises in the material of Khao Ta Phlai, 221 were complete elements (6.9%), and 370 were nearly complete (at least 90% of the complete bone), while 760 (24%) were small fragments representing less than 5% of the complete anatomical part. The average percentage of completion of the bones is 29%. This value is similar in the Neolithic layers of TP1 and TP2 (29%). It is slightly lower in the Metal Ages layer of TP1 (25%), and higher in the same period layers from TP2 (39%), but the small size of this assemblage does not allow considering this result as significant.

The anatomical distributions of the turtle bone elements show strong variations among the different layers (Fig. 6). In the Neolithic layers of TP2, the distribution is fairly homogeneous (mean PR=33%) with representation of all the anatomical parts except for the smallest elements (phalanges, carpal and tarsal articulations, and vertebrae), and the skull. The most robust anatomical parts are the best represented (peripheral plates, epiplastron, entoplastron, and nuchal plate -PR>38%-), while the most fragile elements are the least represented (zeugopods, and most girdle elements -PR<5%-). An exception to this pattern is observed for the peripheral plates of the bridge (PR=15%), which are less represented than the other elements of the carapace and other peripherals (mean PR=73%). The distribution pattern is less homogeneous in TP1, where the mean PR is lower (24% in the Neolithic layers and 15% in the Metal Ages layers), but the Neolithic layers follow the same general pattern as TP2, with a lower global representation due to the strong presence of a single peripheral plate rank. In this last layer, another difference with TP2 is that the stylopods are also better represented than the elements from the carapace. The anatomical distribution of the bones collected in the Metal Ages layers of TP1 is very different, with a very strong representation of the stylopods (mean PR=76%) compared to the most robust elements of the carapace (PR=29-15%). Otherwise, the same general observations apply with a lack of skull and extremity elements, a better representation of the most robust elements of the carapace, and a lower representation of the peripheral plates of the bridge (PR=5%) compared to other peripheral plates (Mean PR=18%).

**Figure 6.**
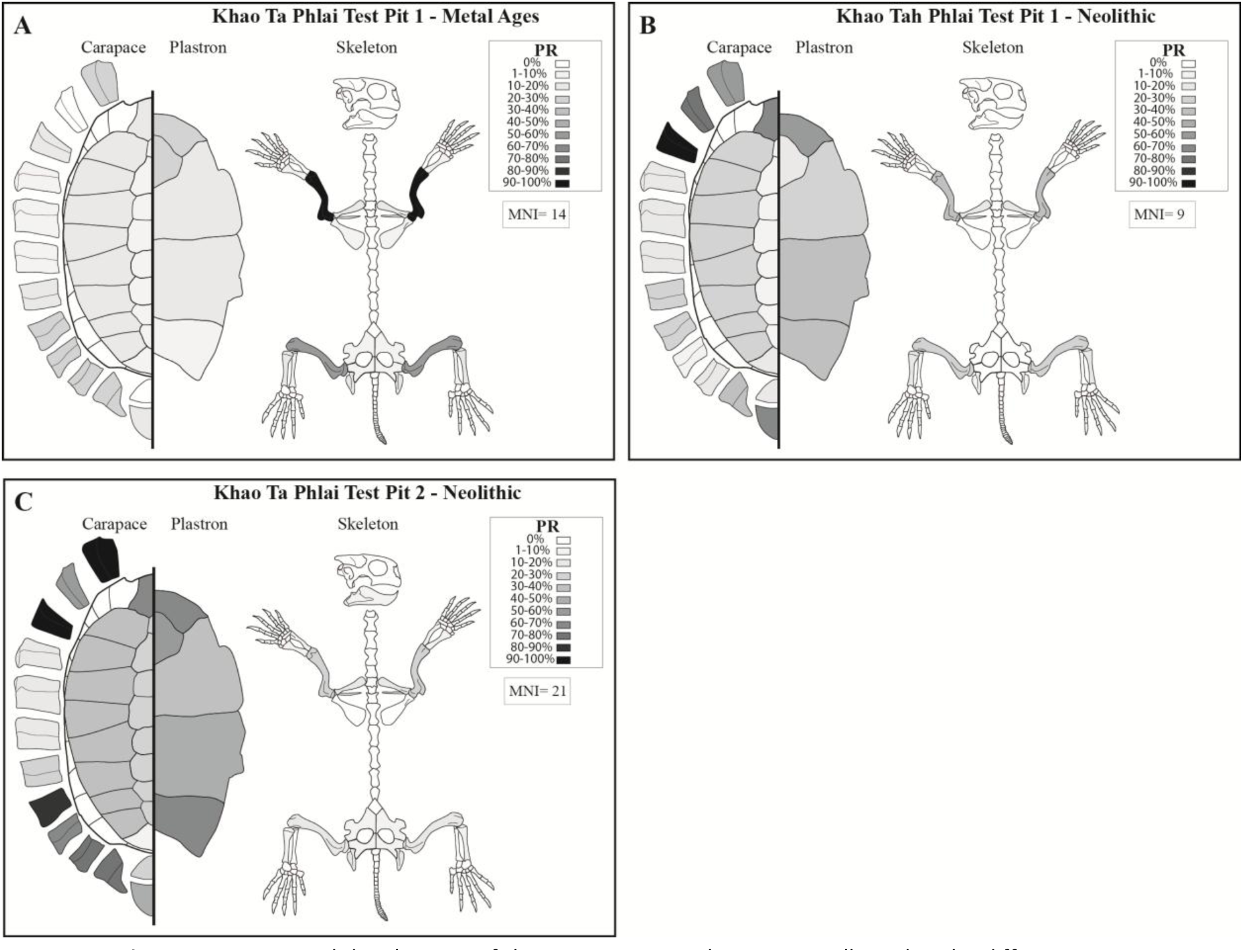
- Anatomical distributions of the non-marine turtle remains collected in the different test-pits and layers of the site of Khao Ta Phlai: (A) Test-pit 1 – Metal Ages, (B) Test-pit 1– Neolithic, (C), Test-pit 2– Neolithic. The percentage of representation (PR) is considered here to provide a graphical visualization of the different values observed for the different anatomical elements.

Most of the material (72% of the NISP) was covered by a veil of calcite, which made the observation of surface alterations of the bones difficult. Despite this limitation, nine bones were recorded as presenting traces of dissolution under the effect of flowing water, and 182 as bearing traces of burning and carbonization. During the study, we also recorded 22 associations of bones from the same context being in anatomical connection in the two TPs below 120 cm of depth in TP2 and 135 cm in TP1. This indicates that the material from the deepest layers was not strongly disturbed since its deposition. Nine combinations of bones in anatomical connection linked together by concretion were also found in the same layers.

##### Size of *Indotestudo elongata* individuals

The measurements recorded on the *I. elongata* archaeological material of the Khao Ta Phlai site have enabled the reconstruction of 219 SCL estimations included between 108 and 252 mm with a mean of 184 mm (Figure. 5-D) and corresponding to at least 26 individuals. Most of the size estimations are from the Neolithic layers of TP2 (n=158) and TP1 (n=31), while only 29 estimations were obtained from the Metal Ages layers (mean size = 178 mm). The strong disparities in the distributions of the size estimations between the archaeological contexts do not allow for an individual comparison of the different layers. The Neolithic layers provided mean SCL values of 185 and 186 mm. The global distribution of these sizes was not unimodal (Hartigans’ dip test, p. value > 0.01), and mixture models indicate it is most likely bimodal with a group of individuals around 210 mm and a second group of smaller specimens around 165 mm.

#### Laang Spean Cave

##### Composition of the herpetofaunal assemblage

The complete herpetofaunal assemblage of Laang Spean consists of 9,533 bone fragments weighing 18,804 grams and representing at least 115 individuals (Tab. 10). Most of them come from the Hoabinhian layer, accounting for 76% of the NISP, 78% of the WR, and 62% of the MNI. The “Neolithic” and “sepulture” assemblages are of similar sizes and account for 10.5% and 13.2% of the NISP, respectively. The material corresponds nearly exclusively to non-marine turtle remains, accounting for 92% of the NISP, 95% of the WR, and 70% of the MNI. Monitor lizards represent 5% of the NISP of the assemblage, snakes are rare (2% of the NISP), and the occurrence of amphibians and smaller lizards is insignificant (below 1%). The distribution of the taxa is not statistically different across the sub-assemblages (Chi² test; p. value > 0.01).

**Table 10.** - Number of Identified Specimens (NISP), Weight of the remains (WR), and Minimum Number of Individuals (MNI) studied in the complete herpetofaunal assemblage from the different layers of the Laang Spean Cave.

|  | Hoabinhian layer |  |  | Neolithic layer |  |  | Sepulture layers |  |  | Total |  |  |
| --- | --- | --- | --- | --- | --- | --- | --- | --- | --- | --- | --- | --- |
|  | NISP | WR | MNI | NISP | WR | MNI | NISP | WR | MNI | NISP | WR | MNI |
| Turtle/Tortoise | 6650 | 13893 | 56 | 904 | 1854 | 12 | 1195 | 2057 | 12 | **8749** | **17804** | **80** |
| Monitor lizard | 406 | 497 | 3 | 30 | 48 | 2 | 39 | 56 | 3 | **475** | **601** | **8** |
| Snake | 158 | 311 | 3 | 16 | 37 | 1 | 19 | 30 | 2 | **193** | **378** | **6** |
| Amphibian | 45 | 9 | 7 | 33 | 6 | 5 | 6 | 1 | 2 | **84** | **16** | **14** |
| Small lizard | 13 | 2 | 3 | 16 | 2 | 3 | 3 | 1 | 1 | **32** | **5** | **7** |
| **Total** | **7272** | **14712** | **72** | **999** | **1947** | **23** | **1262** | **2145** | **20** | **9533** | **18804** | **115** |

The identification of turtle and tortoise bone fragments in Laang Spean Cave (Tab. 11) shows that only 33% of the NISP and 52% of the WR have been attributed to at least a family. *Indotestudo elongata* bone remains account for the majority of the identified turtle/tortoise bones, representing 86% of the NISP, 86% of the WR, and 87% of the MNI. Geoemydidae are rare, with only 14% of the NISP, 13% of the WR, and 9% of the MNI, while the occurrence of Trionychidae is minimal, accounting for less than 1% of the NR and WR, and 3.5% of the MNI.

**Table 11.** - Number of Identified Specimens (NISP), Weight of the remains (WR), and Minimum Number of Individuals (MNI) identified in the turtle/tortoise bones assemblage from the different layers of Laang Spean Cave.

|  | Hoabinhian layer |  |  | Neolithic layer |  |  | Sepulture layers |  |  | Total |  |  |
| --- | --- | --- | --- | --- | --- | --- | --- | --- | --- | --- | --- | --- |
|  | NISP | WR | MNI | NISP | WR | MNI | NISP | WR | MNI | NISP | WR | MNI |
| *Indotestudo elongata* | 2046 | 6706 | 51 | 204 | 649 | 12 | 260 | 692 | 12 | 2510 | 8047 | 75 |
| Geoemydidae | 325 | 955 | 4 | 34 | 115 | 2 | 47 | 162 | 2 | 406 | 1232 | 8 |
| Trionychidae | 7 | 20.5 | 1 | 2 | 7 | 1 | 5 | 16 | 1 | 14 | 43.5 | 3 |
| Turtle ind. | 4272 | 6212 |  | 664 | 1084 |  | 883 | 1187 |  | 5819 | 8483 | 0 |
| **Total** | **6650** | **13894** | **56** | **904** | **1855** | **15** | **1195** | **2057** | **15** | **8749** | **17806** | **86** |

##### Taphonomy of the turtle/tortoise bone assemblage

Regarding the taphonomy of the turtle bones collected in the different squares, the mean completion rate is slightly lower in the “sepulture” assemblage (32%) than in the “Neolithic” and “Hoabinhian” assemblages (39.2% and 36.7%). The general fragmentation pattern is otherwise similar in all layers. The complete bones constitute between 9.3% of the assemblages for the Hoabinhian assemblage and 7.2-6.2% for the “Neolithic” and “Sepulture” assemblages, while nearly complete elements account for 14.2% of the “Hoabinhian”, 13.8% of the “Neolithic”, and 11% of the “Sepulture” assemblages. Most anatomical parts are represented in the different assemblages, but the extremities, vertebrae, and skull remains are very rare with a PR below 4% in all assemblages (Fig. 7). There is also a global tendency towards a lower representation of the peripheral plates of the bridge (11%-31%) in comparison to the other peripheral plates (58%-70%), although this trend is more strongly marked in the Hoabinhian layer (11% vs. 70%). The peripheral plates of the bridge are also systematically more fragmented (59%-67% of mean completion) than the others (83%-87% of mean completion). Outside of these common trends, significant strong differences emerge between the “Hoabinhian” assemblages and the two other layers. Indeed, although the general PR is similar in the different assemblages (41% for the Hoabinhian assemblage, 37% for the Neolithic assemblage, and 34% for the Sepulture assemblage), the carapace elements are dramatically better represented in the Hoabinhian bones (Fig. 7-A) compared to the two other assemblages (Fig. 7-B, C) (Chi² test; p. value < 0.01). Specifically, while the PR of the carapace and plastron bones is more or less similar in the Hoabinhian assemblage (57.8% vs. 67%), the plastron elements are mostly missing in the other assemblages (54% vs. 20.8% in the Neolithic assemblage and 53% vs. 15.1% in the Sepulture assemblage). The stylopods are also better represented in the Hoabinhian squares (84%) compared to the other assemblages (50.4% and 28%).

**Figure 7.**
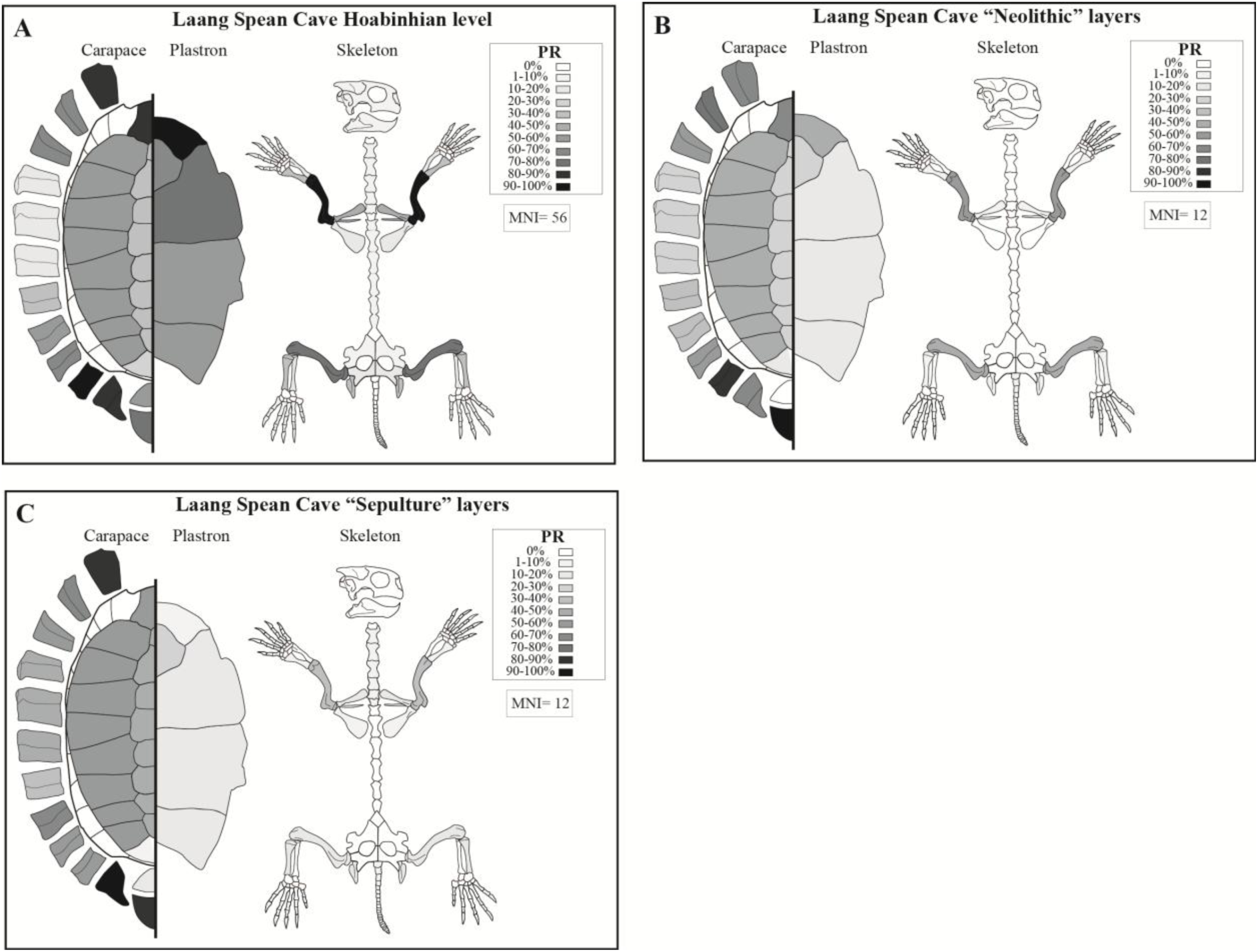
- Anatomical distributions of the turtle remains collected in the different layers of the site of Lang Spean. The percentage of representation (PR) is considered here to provide a graphical visualization of the different values observed for the different anatomical elements.

The observation of traces on the bones is made very challenging by the fact that 61% of them are covered by a veil of calcite. Interestingly, this calcite deposit was more frequent in the Hoabinhian squares, where it covered 65% of the bones, but was scarcer in the Neolithic and sepulture squares, where it covered respectively 56% and 38% of the bones. This is probably related to the position of the remains in the cave, more or less close to the walls, which influenced their exposure to water flows during the rainy season. There is also a possibility that the calcite veil might be more frequent on the oldest remains. The presence of water flow in the site is also indicated by the occurrence of 61 bones that have been polished by water flows.

Porcupine gnawing traces were observed on only 13 elements distributed in several areas and layers of the site, and digestion traces on only one. This clearly indicates a minor impact of animal species on the integrity of the archaeological assemblage. Putative burning traces were observed on 10% of the remains. These traces were better represented in the Hoabinhian squares, where they were present on 12% of the bones, while they only occurred on 2.8% and 3.8% of the bones recovered in the Neolithic and Sepulture squares. The characterization of burning traces was made very difficult by the fact that the material presents a strong variability of surface color, probably related to post-depositional chemical alteration. The occurrence of these traces might thus have been underestimated, given the fact that we chose to record them only when their nature was indisputable. No cut marks were observed.

Among the full assemblage, 327 fragments of carapaces (3.7% of the NR) were still in anatomical connection at the moment of excavation. These elements are mostly from the Hoabinhian squares (N=292), where they account for 4.3% of the turtle remains. Elements in anatomical connection are scarcer in the other assemblages, with only 35 occurrences (1.7% of the turtle NR). This indicates that the Hoabinhian squares have indeed been less disturbed than the “Neolithic” and “Sepulture” squares.

##### Size of *Indotestudo elongata* individuals

The measurements recorded on the *I. elongata* archaeological material of Laang Spean Cave enabled the reconstruction of 688 SCL estimations, ranging from 68 to 345 mm, with a mean of 201 mm (Figure. 5-C), corresponding to at least 75 individuals. Most of the data (N=564) are from Hoabinhian layers, while the Neolithic squares only provided 124 SCL data. However, no significant difference emerged from the comparison of these two assemblages (Student t-test, p. value > 0.05). The global distribution (across all squares) of these sizes is unimodal (Hartigans’ dip test, p. value > 0.05) with a peak of specimens around 200 mm SCL. In this site, small specimens below 170 mm represent only 16% of the population, and specimens below 140 mm only constitute 4.7%.

## Discussion

### Taxonomic composition of the herpetofaunal assemblages

In all the assemblages, the distribution of herpetofaunal groups in the four investigated sites shows strong similarities. Non-marine turtles are nearly always the best-represented herpetofaunal group (between 59 and 91% of the NISP), followed by Monitor lizards (between 6 and 25% of the NISP), snakes (below 3.5% of the NISP), and amphibians. The only exception to this trend is layer 1 of Moh Khiew cave, where bone remains of Monitor lizards (25% of the NISP) and snakes (23% of the NISP) are more numerous than turtle skeletal elements (20% of the NISP). However, this layer is disturbed and not dated, making interpretation impossible at this time. Nonetheless, the other sites follow a clear pattern, indicating that hunter-gatherer groups have preferred to exploit turtles over other reptile and amphibian taxa, in line with previously observed regional patterns in similar zooarchaeological assemblages (Conrad, 2015).

Regarding the proportion of turtle/tortoise families in the assemblages, Testudinidae (*Indotestudo elongata*) always represents the most significant group, accounting for between 52% and 89% of the turtle bone NISPs. The proportions of Geoemydidae turtles vary widely, ranging from 48% to 11% of the same NISPs. The variability could be explained by the accessibility of streams, rivers, and lakes by the inhabitants of the sites, as most of the species from this group are aquatic freshwater turtles. Generally, Geoemydidae accounts for around 30% of the NISP in most sites, but they are less prevalent in Laang Spean Cave and are best represented in the TP1 of the Khao Ta Phlai site. Data from the faunal assemblage of Laang Spean Cave also indicate a weak contribution of freshwater taxa (mussels and fish) to the overall diet (Forestier et al., 2015; Frère et al., 2018), which aligns with the observation of scarcity of freshwater turtles in the site. Regarding the prevalence of Geoemydidae species in the TP1 of Khao Ta Phlai, it might suggest a stronger reliance on freshwater resources compared to other sites. However, considering that the chronology of the two test-pits of the site is not yet fully resolved and that the general importance of aquatic resources in this assemblage needs further estimation, this fact cannot be related to a cultural/chronological trend at this stage. Overall, the data on herpetofaunal assemblages point to strong similarities between assemblages of different ages and from various environmental settings. This aspect should be considered in conjunction with studies on mammal bone assemblages of the same sites to test the hypothesis of a potential homogeneity of Hoabinhian subsistence strategies in continental Southeast Asia.

### Taphonomy of the turtle assemblages

The fragmentation rate of the bones is fairly consistent among the sites, with an average percentage of completeness ranging from 37% to 28%. The material from the first three layers of Moh Khiew Cave and Laang Spean Cave shows the least fragmentation, with an average percentage of completeness above 33%. However, the layer 4 of Moh Khiew cave provided the most fragmented material, with an average percentage of completeness of 28%. The presence of large limestone blocks in this layer might indicate crumbling that could have altered the faunal material.

Regarding the anatomical distribution of the turtle remains, the sites present significant differences, with mean PR between 41% (Laang Spean Cave) and 15% (Khao Ta Phlai Metal Ages layer from TP1). This means that the anatomical representation of the bone remains is more or less strongly biased towards certain elements. Two main scenarios occur in the assemblages: sites where stylopods are the best-represented parts (Khao Ta Phlai Metal Ages layer from TP1, Doi Pha Kan site, layers 2 to 4 of Moh Khiew cave), and sites where the most robust parts of the carapace are the best-represented elements. The mean PR is systematically higher in the assemblages where the stylopods are most numerous, indicating that these assemblages are less altered by post-depositional phenomena. Indeed, a natural alteration would rather lead to the situation observed in the other assemblages, in which the most robust elements are the most frequently found, and thus, they would have the highest survival rates. However, this is not sufficient to explain an overrepresentation of long bones, which are supposed to preserve less well than carapace elements. Considering that all the sediment of the studied deposit has been screened, a major recovery bias is unlikely, although some of the smallest elements might have been missed. A post-depositional sorting of the material could also be ruled out, as we have shown no evidence of differential fragmentation and no abundant traces of water circulation in the different deposits studied. The most likely hypothesis is thus that human inhabitants of some sites discarded or transported some carapaces of consumed animals for further use and left the smallest elements, among which the largest and toughest (humerus and femurs), have been recovered and identified. This behavior, however, does not seem to be systematic, as the anatomical distributions indicate that complete individuals have been brought to the sites. The absence of the head of the specimens could be either related to an identification bias or a removal of these parts outside of the site. Apart from the humerus and femur, the anatomical distributions of turtle bones follow a global pattern where the most robust anatomical elements are better represented than the more fragile ones. The peripheral plates of the bridge are an exception to this trend, always being less represented than the other peripherals. This is likely related to an identification bias due to the nearly complete absence of complete pieces of such elements in the material, likely linked to the separation of the carapace from the plastron by the inhabitants of the sites who broke the bones in the area that links both parts of the shell to access most of the meat content of the animal.

The observations of surface traces on the bones indicate a nearly complete lack of predation and digestion traces. Combined with the general low fragmentation of the material, this completely rules out a significant role of non-human predators in the formation of the studied assemblages. This is consistent with the fact that, although some predators, including Monitor lizards, are known to hunt juvenile tortoise individuals, adult tortoises likely have few non-human predators, although some modern specimens bear traces of predation attempts (Ihlow et al., 2016), and predation on other Southeast Asian tortoise species has been reported (Platt et al., 2021). Large felids (Emmons, 1989) and eagles (Gil-Sánchez et al., 2022) are known to be able to hunt adult tortoises, but such predators would undoubtedly leave predation traces on the subfossil bone assemblages studied. Some very rare bones bearing porcupine traces indicate that these animals had a minor impact on some of the assemblages, but not enough to impact the zooarchaeological interpretations. However, although it seems fairly evident that the animals present in the sites have been hunted to be consumed as there is no trace of bone industry in the assemblages, finding direct traces of culinary preparation on the bones is very challenging. In Khao Ta Phlai and Laang Spean, some remains (72% and 37%) were covered by a veil of calcite, making it impossible to observe the bone surfaces. Additionally, very few cut marks have been identified on the bones from the different sites. Many burned bones were observed in all the sites, but linking these to cooking techniques is questionable. These traces do not seem to be located on specific parts of the bones (e.g., the external side of the carapace) and appear randomly on every area of every anatomical part. It is likely that these traces are related to post-depositional events unrelated to the cooking of the animals. The frequency of fire traces, combined with the strong fragmentation of large vertebrate remains in most sites (C. Griggo; C. Bochaton pers. obs), could indicate the use of bones as fuel (Villa et al., 2002). Such a use is unlikely for turtle skeletons considering their small size and the very good preservation stage of their remains, but proximity to fireplaces (Bennett, 1999) could explain the random occurrence of fire traces on their bone elements.

The similarities observed among the turtle assemblages at these sites could potentially suggest a shared approach to the management of turtle carcasses by hunter-gatherers and/or uniformity in the overall use of the studied sites. It is important to note, however, that a comprehensive discussion of these possibilities is constrained by the absence of complete zooarchaeological studies, lithic analyses, and geoarchaeological observations across all the deposits under investigation. Furthermore, it is crucial to recognize that such taphonomic homogeneity does not necessarily imply cultural similarities, as similar carcass management practices can be employed by disparate human groups with distinct cultural contexts.

### Size of Indotestudo elongata archaeological specimens

The size of *I. elongata* individuals observed in the four archaeological deposits (Fig. 5) shows common patterns but also some differences. The distributions of estimated sizes are bimodal in all sites except Laang Spean. In all the sites, most of the estimations correspond to adult-size specimens above 170 mm SCL, reaching maximums of 270-345 mm SCL. These specimens fall within the size range of modern representatives of the species. However, all sites present a variable proportion of smaller, likely immature individuals. The representation of this second group is the lowest in Laang Spean (16% of the total number of estimations) but is significant enough in the other sites to make their distributions bimodal, with 35%-33% in Doi Pha Kan and Khao Ta Phlai, and 24% in Moh Khiew cave. Specimens below 140mm SCL are rare in all sites, accounting for less than 10% of the estimations in all sites, but represent more than 15% of the Doi Pha Kan population.

Interpreting the size distribution of the archaeological tortoises is a challenging task as it requires an idea of what the size structure of a wild population would look like, along with basic biological data (e.g., season of birth, activity pattern, growth speed) regarding modern and past *I. elongata* populations. However, this data is mostly missing, making a detailed interpretation of the collected archaeological data challenging. The recovery of size distribution data in a natural modern population is always challenging as it could be influenced by various factors (e.g., climate, environment, seasonality, behaviors, and sizes of the individuals) that could bias the observations and make some size classes more difficult to observe than others. Moreover, the history and specific conditions of a wild population itself could have a dramatic impact on its size structure, further complicating the comparison with archaeological populations.

To our knowledge, the only data collected on *I. elongata* concern the population of the Ban Kok Village (Khon Kaen Province, Thailand). This study shows that the pre-adult individuals have a low survivability rate, as their population mostly consists of newly born and old adult individuals (Sriprateep et al., 2013). The authors suggest that this strongly biased structure could be related to an absence of predation on the large individuals and partly to several phenomena having a stronger impact on small specimens (e.g., predation, trampling of domestic bovids). However, the main cause is still unknown, and a potential poaching of the smaller individuals is not discussed. Similarly, another publication about *I. travancorica* indicates a lack of juvenile specimens in the population but highlights that it could be related to a seasonal activity-specific pattern. Juvenile specimens were much more commonly found at the beginning of the rainy season than during the dry season when their study was conducted (Ramesh, 2008). Other published distributions from other tortoise species also indicate a strong representation of adult-size individuals of different ages having completed their growth, but they also show a more balanced distribution of juvenile specimens of all sizes (Hailey and Coulson, 1999; Rouag et al., 2007; Znari et al., 2005). In all these distributions, the juvenile specimens are scarcer than adult ones, which makes sense as adult-class specimens correspond to individuals of very different ages that have reached their final size. The only case in which this situation would be reversed is a population in which adult individuals would be subject to a strong predation pressure superior to the pressures imposed on the smaller individuals.

The site of Laang Spean Cave presents a unimodal size distribution in which juvenile specimens are mostly excluded. In that sense, this distribution is very different from that of a natural population and indicates a strong selection on adult specimens of moderate to large size. This is clearly indicative of a very selective hunting strategy that may have been enabled by the abundance of resources in the vicinity of the site. Such a selection, although visible in other deposits, is less marked as juvenile specimens compose a more significant part of the assemblages, especially in Doi Pha Kan. In these sites, it is impossible to estimate whether or not the proportions of juvenile specimens present in the assemblages are similar to those of the exploited natural populations and thus to estimate the exact intensity of the selection toward large size individuals. In any case, it is the sign of an opportunistic foraging strategy, as such a combination of juvenile specimens has been observed in modern hunter-gatherer populations actively collecting tortoises this way (Mena et al., 2000). However, this might also be influenced by the hunting method, in the case a direct selection by the hunter is not made, for instance with the use of trapping that was also hypothesized in Doi Pha Kan for the hunting of monitor lizards (Bochaton et al., 2019a). This technique is also the most used to hunt tortoises in the Amazon, as it is the most efficient method before active searching (Santos et al., 2020). This implies no selection on the specimens in the wild, although the type of trap used (e.g., size of the ground hole) might induce some size bias. The use of traps could thus explain the strong representation of smaller individuals present in the archaeological assemblages and indicate a very opportunistic strategy, indicative of either a poor selection by the hunter and/or a relative scarcity of the tortoises in the environments, making it harder to collect large individuals. The hunting season could also be an explanation for the stronger or weaker presence of juvenile specimens in the assemblages. During the dry season, tortoises are less active and harder to find, which could lead the hunter to be less selective, especially in the case of the use of non-selective hunting methods that allow them to find these animals. Theobald (1868) mentions the hunts of tortoises by Burmese hunters in the dry season by clearing grasslands and forests with fire to destroy their shelters and locate them. In contrast, smaller tortoises are more active in the rainy season, during which dogs are more used to track them (Blythe, 1854; Theobald, 1868). Ultimately, both seasonal hypotheses could explain the occurrence of small individuals, using different explanations (hunting method vs. activity season). Only the use of non-traditional approaches, such as skeletochronology (Ehret, 2007), could help to clarify this question by estimating the season of death of the tortoise individuals, as well as the occupation seasonality of the different sites, given the absence of other seasonality markers in the materials.

Tortoise populations are vulnerable to intensive exploitation, often targeting larger mature individuals. Consequently, their exploitation has been viewed as an indicator of small-scale hunting and, thus, of relatively small human groups (Stiner et al., 2000). In the sites under study, the pronounced focus on a single turtle species (*I. elongata*) and the emphasis placed on larger individuals could potentially lead to detrimental consequences for natural populations. This could involve a sustained reduction in the number of individuals and a decrease in average specimen size over the long term (Close and Seigel, 1997). Such exploitation could remain sustainable only if it were not intense, implying that a relatively limited number of individuals were harvested to sustain a potentially small-sized human group. Evaluating this aspect proves challenging, as comprehending the overall significance of tortoises in the diet of Southeast Asian hunter-gatherer groups studied, and thus estimating the intensity of their exploitation, requires a comprehensive and quantified examination of the mammal fauna at the sites, as well as robust data pertaining to occupation duration and site usage. Nonetheless, it is evident that the prehistoric populations under investigation did engage in the exploitation of tortoises, which constituted a notable component of their meat-based diet. This is not surprising, as turtle species are supposed to represent an important biomass in the ecosystems (Iverson, 1982) and are also fairly easy to collect. This behavior has persisted until nowadays in continental Southeast Asia hunter-gatherer modern groups (Hansel, 2004), although not all populations choose to exploit reptile species (Tungittiplakornl and Dearden, 2002).

## Conclusion and Perspectives

This work has been developed as a foundation, aiming to furnish fundamental data and research instruments essential for investigating tortoise assemblages in continental Southeast Asia. Consequently, the full extent of this effort’s value will be realized by employing its analytical methodology in forthcoming studies and juxtaposing it against supplementary assemblages for comparison. We were, however, able to reach several conclusions, as we demonstrated potential strong similarities between the exploitation of herpetofaunal taxa in the different sites, as well as in the taphonomy of the non-marine turtle assemblages in different chronological and environmental settings. These data thus open many interesting questions regarding the trends of hunter-gatherer subsistence strategies in continental Southeast Asia during the Pleistocene and through the Holocene. However, much work remains to be done to reach a satisfactory zooarchaeological documentation level regarding these prehistoric human groups.

As we demonstrated it in the introduction of this paper, many of the previously excavated Hoabinhian archaeological deposits of continental Southeast Asia, including sites which are known to have provided rich assemblages of non-marine turtle bone remains (e.g., Lang Rongrien), have not benefited from quantified zooarchaeological analyses. The complete study of these sites will be important to provide additional relevant comparison points to the present study. The non-herpetofaunal taxa of the sites included in this study should also be investigated to estimate the relative part of the reptile and amphibian exploitation in the global diet of these hunter-gatherer populations. Such studies should be carried out in combination with the elaboration of appropriate study protocols regarding the estimation of the size/weight of the exploited individuals of large mammal species. Much-needed is also the elaboration of identification methods, whether morphological or molecular, designed for Southeast Asian species to complement the existing works (Bochaton et al., 2019b; Pritchard et al., 2009). Only at the cost of such investment will the zooarchaeology of Southeast Asia be on par with the rich literature existing on the material productions of prehistoric groups. However, this will also require the local development of a strong research community interested in that discipline, which is still lacking at the present.

## Acknowledgments

The authors are grateful to all the excavation teams who collected the studied remains, as well as to all the technical staff who helped us in the study of the different bone assemblages. We are especially grateful to Silpakorn University, which hosted C. B. during most of the studies, as well as to the Fine Art Department of Thailand. We also thank S. Coleman and the Florida Museum of Natural History for welcoming us in their herpetology collections. We are also grateful to the PCI Archaeology recommender who handled this work (Ruth Blasco) and to the two reviewers who helped us to improve this manuscript (Iratxe Boneta and Noel Amano).

## Data, scripts, code, and supplementary information availability

Supplementary information is available online: https://doi.org/10.1101/2023.04.27.538552

## Conflict of interest disclosure

The authors declare that they comply with the PCI rule of having no financial conflicts of interest in relation to the content of the article.

## Funding

This work has been done thanks to the support of several funding agencies: the FYSSEN foundation, the DIM-MAP funded by the Region île-de-France, the French Ministry of Europe and Foreign Affairs, and the IRN PalBioDivASE 0846 funded by the CNRS. It was also supported by the Mission Préhistorique Franco-Cambodgienne, the Mission Préhistorique Franco-Thaïe, and the Mission Paléolithique Franco-Thaïe of the French Ministry of Europe and Foreign Affairs (MEAE, Paris).

## Appendix

### Appendix 1: All measurements recorded on the modern turtle skeletons

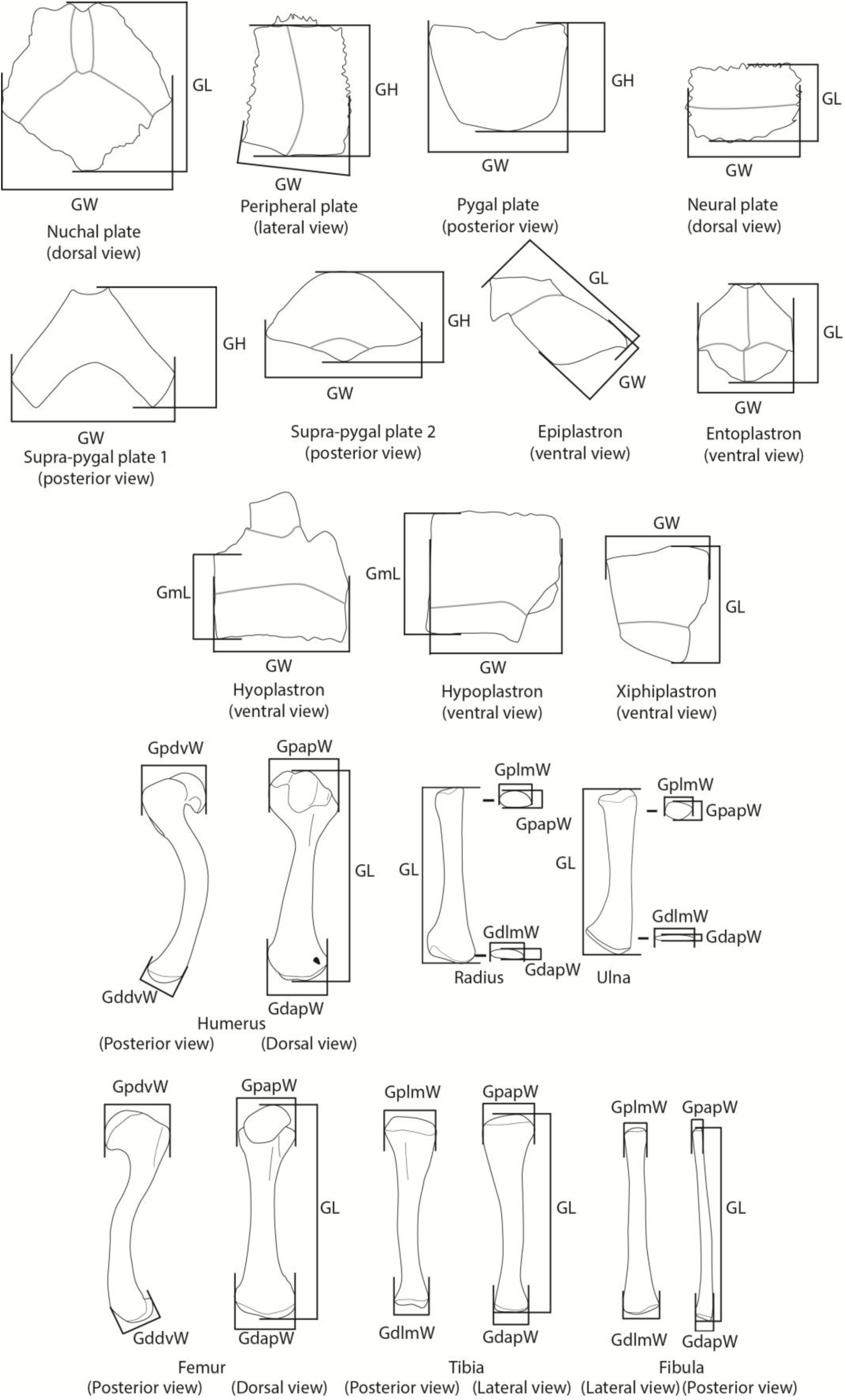

## Notes

### Competing Interest Statement

The authors have declared no competing interest.

### Summary of Updates

Adding of the recommendation logo of PCI Archaeology; Removal of a duplicated sentence in the abstract and of the line numbering; author affiliations updated

## References

1. Adi, B.T.H., 2000. Archaeology of Ulu Kelantan (PhD thesis). Australian National University, Canberra.

2. Amphansri, A., 2011. Temporal and Spatial Analysis of Faunal Remains from Tham Lod Rockshelter Site, Pang Mapha District, Mae Hong Son Province (M. A. Thesis). Silpakorn University, Bangkok, Thailand.

3. Anderson, D.D., 1990. Lang Rongrien Rockshelter: A Pleistocene, Early Holocene Archaeological Site from Krabi, Southwestern Thailand. University of Pennsylvania Press, Philadelphia.

4. Auetrakulvit, P., 2004. Faunes du pléistoc ne nal l’holoc ne de Tha lande : approche archéozoologique (PhD thesis). Aix Marseille 1, Marseille.

5. Auetrakulvit, P., Forestier, H., Khaokhiew, C., Zeitoun, V., 2012. New excavation at MohKhiew site (Southern Thailand), in: Bonatz, D., Reinecke, A., Tjoa-Bonatz, M.L. (Eds.), Crossing Borders in Southeast Asian Archaeology. Singapore, pp. 62–74.

6. Auffenberg, W., 1974. Checklist of Fossil Land Tortoises (Testudinidae). Bulletin of the Florida State Museum Biological Sciences 18, 122–247.

7. Avery, G., Kandel, A.W., Klein, R.G., Conard, N.J., Cruz-Uribe, K., 2004. Tortoises as food and taphonomic elements in palaeo « landscapes », in: Petits Animaux et Sociétés Humaines: Du Complément Alimentaire Aux Ressources Utilitaires. Presented at the XXIVe Rencontres internationales d’archéologie et d’histoire d’Antibes, Antibes, France, pp. 147–161.

8. Bennett, J.L., 1999. Thermal Alteration of Buried Bone. Journal of Archaeological Science 26, 1–8. 10.1006/jasc.1998.0283

9. Biton, R., Sharon, G., Oron, M., Steiner, T., Rabinovich, R., 2017. Freshwater turtle or tortoise? The exploitation of testudines at the Mousterian site of Nahal Mahanayeem Outlet, Hula Valley, Israel. Journal of Archaeological Science: Reports 14, 409–419. 10.1016/j.jasrep.2017.05.058

10. Blasco, R., 2008. Human consumption of tortoises at Level IV of Bolomor Cave (Valencia, Spain). Journal of Archaeological Science 35, 2839–2848. 10.1016/j.jas.2008.05.013

11. Blasco, R., Rosell, J., Smith, K.T., Maul, L.C., Sañudo, P., Barkai, R., Gopher, A., 2016. Tortoises as a dietary supplement: A view from the Middle Pleistocene site of Qesem Cave, Israel. Quaternary Science Reviews 133, 165–182. 10.1016/j.quascirev.2015.12.006

12. Blythe, E., 1854. Notices and descriptions of various reptiles, new or little-known. Part I. The journal of the Asiatic Society of Bengal 22, 639–655.

13. Bochaton, C., Hanot, P., Frère, S., Claude, J., Naksri, W., Auetrakulvit, P., Zeitoun, V., 2019a. Size and weight estimations of subfossil monitor lizards (Varanus sp. Merrem 1820) with an application to the Hoabinhian assemblage of Doi Pha Kan (Late Pleistocene, Lampang province, Thailand). Annales de Paléontologie 4, 295–304. 10.1016/j.annpal.2019.05.003

14. Bochaton, C., Ivanov, M., Claude, J., 2019b. Osteological criteria for the specific identification of Monitor lizards (*Varanus* Merrem, 1820) remains in subfossil deposits of Sundaland and continental Southeast Asia. Amphibia-Reptilia 40, 219–232. 10.1163/15685381-20181101

15. Bulbeck, F., 2003. Hunter-Gatherer Occupation of the Malay Peninsula from the Ice Age to the Iron Age, in: Under the Canopy: The Archaeology of Tropical Rain Forests. Rutgers University Press, Brunswick, pp. 119–160.

16. Celiberti, V., Forestier, H., Auetrakulvit, P., Zeitoun, V., 2018. Doi Pha Kan, Lampang Province, North of Thailand: A New Lithic Assemblage in the Hoabinhian Chrono-Cultural Context, in: Tan, N. (Ed.), Advancing Southeast Asian Archaeology. Bangkok, pp. 91–99, 316–319.

17. Chung, T.N., 2008. Research on Hoa Binh culture in Viet Nam, Laos and Cambodia. Vietnam Archaeology 3, 19–32.

18. Claude, J., Auetrakulvit, P., Naksri, W., Bochaton, C., Zeitoun, V., Tong, H., 2019. The recent fossil turtle record of the central plain of Thailand reveals local extinctions. Annales de Paléontologie 4, 305–315. 10.1016/j.annpal.2019.04.005

19. Close, L.M., Seigel, R., 1997. Differences in body size among populations of red-eared sliders (Trachemys scripta elegans) subjected to different levels of harvesting. Chelonian Conservation and Biology 2, 563–566.

20. Codron, D., Holt, S., Wilson, B., Horwitz, L.K., 2022. Skeletal allometries in the leopard tortoise (Stigmochelys pardalis): Predicting chelonian body size and mass distributions in archaeozoological assemblages. Quaternary International, Quaternary Environments and Archaeology of the Northern Cape (South Africa) 614, 59–72. 10.1016/j.quaint.2021.07.021

21. Collectif, 1932. Premier congr s des préhistoriens d’Extrême-Orient. Imprimerie d’Extrême-Orient, Hanoi.

22. Conrad, C., 2015. Archaeozoology in Mainland Southeast Asia: Changing Methodology and Pleistocene to Holocene Forager Subsistence Patterns in Thailand and Peninsular Malaysia. Open Quaternary 1. 10.5334/oq.af

23. Conrad, C., Higham, C., Eda, M., Marwick, B., 2016. Palaeoecology and Forager Subsistence Strategies during the Pleistocene – Holocene Transition: A Reinvestigation of the Zooarchaeological Assemblage from Spirit Cave, Mae Hong Son Province, Thailand. Asian Perspectives 55, 2–27. 10.1353/asi.2016.0013

24. Das, I., 2010. A field guide to the reptiles of South-East Asia. Bloomsbury Publishing, London, New Delhi, New York, Sydney.

25. Dodson, P., Wexlar, D., 1979. Taphonomic Investigations of Owl Pellets. Paleobiology 5, 275–284.

26. Dunn, F.L., 1964. Excavations at Gua Kechil, Pahang. Journal of the Malaysian Branch of the Royal Asiatic Society 37, 87–124.

27. Eberling, G., 2011. Haltung und Nachzucht von *Indotestudo elongata* (Blythe, 1853). Draco 2, 61–72.

28. Ehret, D., 2007. Skeletochronology: A Method for Determing the Individual Age and Growth of Modern and Fossil Tortoises (Reptilia: Testudines). Bulletin of the Florida Museum of Natural History 47, 49–72.

29. Emmons, L.H., 1989. Jaguar Predation on Chelonians. Journal of Herpetology 23, 311–314. 10.2307/1564460

30. Esker, D.A., Forman, S.L., Butler, D.K., 2019. Reconstructing the mass and thermal ecology of North American Pleistocene tortoises. Paleobiology 45, 363–377. 10.1017/pab.2019.6

31. Fernàndez-Jalvo, Y., Andrews, P., 2016. Atlas of Taphonomic Identifications: 1001+ Images of Fossil and Recent Mammal Bone Modification. Springer, London, UK.

32. Forestier, H., 2003. Des outils nés de la forêt : l’importance du végétal en Asie du Sud-Est dans l’imagination et l’invention technique aux périodes préhistoriques, in: Froment, A., Guffroy, J., Peuplements Anciens et Actuels des Forêts Tropicales : Séminaire-Atelier, Orléans (FRA), 1998/10/15-16 (Eds.), Peuplements anciens et actuels des forêts tropicales : actes du séminaire-atelier, Colloques et Séminaires. IRD, Paris, pp. 315–337.

33. Forestier, H., Griggo, C., Sophady, H., 2023. L’Hoabinhien ou le paradigme égaré de la modernité européenne en Extrême-Orient : l’exemple de la Préhistoire du Cambodge, in: Le Poids de l’histoire Des Sciences et l’hégémonie Européenne En Préhistoire. Presented at the 29e Congr s préhistorique de France, Société préhistorique française, Toulouse, pp. 59–76.

34. Forestier, H., Sophady, H., Celiberti, V., 2017. Le techno-complexe hoabinhien en Asie du Sud-est continentale : L’histoire d’un galet qui cache la forêt. Journal of Lithic Studies 4, 305–349. 10.2218/jls.v4i2.2545

35. Forestier, H., Sophady, H., Puaud, S., Celiberti, V., Frère, S., Zeitoun, V., Mourer-Chauviré, C., Mourer, R., Than, H., Billault, L., 2015. The Hoabinhian from Laang Spean Cave in its stratigraphic, chronological, typo-technological and environmental context (Cambodia, Battambang province). Journal of Archaeological Science: Reports 3, 194–206. 10.1016/j.jasrep.2015.06.008

36. Forestier, H., Zeitoun, V., Winayalai, C., Métais, C., 2013. The open-air site of Huai Hin (Northwestern Thailand): Chronological perspectives for the Hoabinhian. Comptes Rendus Palevol 12, 45–55. 10.1016/j.crpv.2012.09.003

37. Forestier, H., Zhou, Y., Auetrakulvit, P., Khaokhiew, C., Li, Y., Ji, X., Zeitoun, V., 2021. Hoabinhian variability in Mainland Southeast Asia revisited: The lithic assemblage of Moh Khiew Cave, Southwestern Thailand. Archaeological Research in Asia 25, 100236. 10.1016/j.ara.2020.100236

38. Forestier, H., Zhou, Y., Viallet, C., Auetrakulvit, P., Li, Y., Sophady, H., 2022. Reduction Sequences During the Hoabinhian Technocomplex in Cambodia and Thailand: A New Knapping Strategy in Southeast Asia from the Terminal Upper Pleistocene to mid Holocene. Lithic Technology 47, 147–170. 10.1080/01977261.2021.1981654

39. Frère, S., Auetrakulvit, P., Zeitoun, V., Sophady, H., Forestier, H., 2018. Doi Pha Kan (Thailand), Ban Tha Si (Thailand) and Laang Spean (Cambodia) Late Paleolithic animal bone assemblages. A new perception of meat supply strategies for Early Holocene Mainland South-east Asia?, in: Tan, N.H. (Ed.), Advancing Southeast Asian Archaeology. Bangkok, pp. 100–108, 320–323.

40. Garbin, R.C., Ascarrunz, E., Joyce, W.G., 2018. Polymorphic characters in the reconstruction of the phylogeny of geoemydid turtles. Zoological Journal of the Linnean Society 184, 896–918. 10.1093/zoolinnean/zlx106

41. Gil-Sánchez, J.M., Rodríguez-Caro, R.C., Moleón, M., Martínez-Pastor, M.C., León-Ortega, M., Eguía, S., Graciá, E., Botella, F., Sánchez-Zapata, J.A., Martínez-Fernández, J., Esteve-Selma, M.A., Giménez, A., 2022. Predation impact on threatened spur-thighed tortoises by golden eagles when main prey is scarce. Sci Rep 12, 17843. 10.1038/s41598-022-22288-9

42. Glover, I.C., 1977. The Hoabinhian hunter-gatherers or early agriculture list in Southeast Asia, in: Megaw, J.V.S. (Ed.), Hunter Gatherers and First Farmer beyond Europe. Leicester University Press, Leicester, UK, pp. 145–166.

43. Gorman, C., 1971. The Hoabinhian and After: Subsistence Patterns in Southeast Asia during the Late Pleistocene and Early Recent Periods. World Archaeology 2, 300–320.

44. Gorman, C.F., 1970. Excavations at Spirit Cave, North Thailand: SOME INTERIM INTERPRETATIONS. Asian Perspectives 13, 79–107.

45. Gorman, C.F., 1969. Hoabinhian: A Pebble-Tool Complex with Early Plant Associations in Southeast Asia. Science 163, 671–673. 10.1126/science.163.3868.671

46. Gould, S.J., 1966. Allometry and size in ontogeny and phylogeny. Biological Reviews 41, 587–640.

47. Guerin, C., Mourer-Chauviré, C., 1969. Le Rhinoceros sondaicus Desmarest du gisement néolithique de Loang Spean, province de Battambang, Cambodge. Ann. Fac. Sc. Phnom Penh, 1970 2, 261–274.

48. Hailey, A., Coulson, I., 1999. The growth pattern of the African tortoise Geochelone pardalis and other chelonians. Canadian Journal of Zoology 77, 181–193. 10.1139/cjz-77-2-181

49. Hansel, T., 2004. Observations on subsistence hunting along the Phu Yai Mountain Range, Xanakham District, Vientiane Province, Lao PDR. Natural History Bulletin of the Siam Society 52, 195–200.

50. Higham, C., 2002. Early Cultures of Mainland Southeast Asia. Art Media Resources.

51. Higham, C.F.W., 1977. Economic Change in Prehistoric Thailand, in: Reed, C. (Ed.), Origins of Agriculture. De Gruyter Mouton, The Hauge, pp. 385–412. 10.1515/9783110813487.385

52. Huxley, J.S., 1932. Problems of relative growth. Methuen & co, London.

53. Ihlow, F., Dawson, J.E., Hartmann, T., Som, S., 2016. *Indotestudo elongata* (Blyth 1854) – Elongated Tortoise, Yellow-headed Tortoise, Yellow Tortoise. Chelonian Conservation and Biology 5, 1–14. 10.3854/crm.5.096.elongata.v1.2016

54. Ihlow, F., Handschuh, M., 2011. Auswilderung von Indotestudo elongata im Kulen Promtep Wildlife Sanctuary im Norden Kambodschas. Marginata 8, 16–23.

55. Imdirakphol, S., Zazzo, A., Auetrakulvit, P., Tiamtinkrit, C., Pierret, A., Forestier, H., Zeitoun, V., 2017. The perforated stones of the Doi Pha Kan burials (Northern Thailand): A Mesolithic singularity? Comptes Rendus Palevol 16, 351–361. 10.1016/j.crpv.2016.12.003

56. Iverson, J.B., 1982. Biomass in turtle populations: A neglected subject. Oecologia 55, 69–76. 10.1007/BF00386720

57. Iverson, J.B., Spinks, P.Q., Shaffer, B.H., McCord, W.P., Das, I., 2001. Phylogenetic relationships among the asian tortoises of the genus *Indotestudo* (Reptilia: Testudines:testudinidae). Hamadryad 26, 283– 286.

58. Ji, X., Kuman, K., Clarke, R.J., Forestier, H., Li, Y., Ma, J., Qiu, K., Li, H., Wu, Y., 2016. The oldest Hoabinhian technocomplex in Asia (43.5 ka) at Xiaodong rockshelter, Yunnan Province, southwest China. Quaternary International, Peking Man and related studies 400, 166–174. 10.1016/j.quaint.2015.09.080

59. Klein, R.G., Cruz-Uribe, K., 1983. Stone Age Population Numbers and Average Tortoise Size at Byneskranskop Cave 1 and DieKelders Cave 1, Southern Cape Province, South Africa. The South African Archaeological Bulletin 38, 26–30.

60. Lyman, R.L., 2008. Quantitative paleozoology, Cambridge University Press. ed, Manuals in archaeology. Cambridge University Press, New-York, USA.

61. Masojc, M., Le, H.D., Gralak, T., Michalec, G., Apolinarska, K., Badura, M., Cendrowska, M., Galas, A., Krupa-Kurzynowska, J., Miazga, B., Ospinska, M., Rozok, Z., Viet, N., 2023. The Early Holocene Hoabinhian (8300-8000 cal BC) occupation from Hiem Cave, Vietnam. Comptes Rendus Palevol 22, 59–76.

62. Medway, Lord, 1969. Excavations At Gua Kechil, Pahang III. Animal Remains. Journal of the Malaysian Branch of the Royal Asiatic Society 42, 197–205.

63. Mena, P., Stallings, J.R., Regalado, J., Cueva, R., 2000. The Sustainability of Current Hunting Practices by the Huaorani. In Hunting for Sustainability in Tropical Forests 57–78.

64. Moser, J., 2001. Hoabinhian: Geographie und Chronologie eines steinzeitlichen Technokomplexes in Südostasien, AVA - Forschungen. Linden Soft, Germany.

65. Mourer-Chauviré, C., Mourer, R., 1970. The Prehistoric Industry of Laang Spean, Province of Battambang, Cambodia. Archaeology and Physical Anthropology in Oceania 5, 128–146. 10.1002/j.1834-4453.1970.tb00110.x

66. Mourer-Chauviré, C., Mourer, R., Thommeret, Y., 1970. Premi res datations absolues de l’habitat préhistorique de la grotte de Laang Spean, province of Battambang (Cambodge). C. R. Acad. Sci. Paris, Ser. D 270, 471–473.

67. Mudar, K., Anderson, D.D., 2007. New Evidence for Southeast Asian Pleistocene Foraging Economies: Faunal Remains from the Early Levels of Lang Rongrien Rockshelter, Krabi, Thailand. Asian Perspectives 46, 298–334.

68. Nabais, M., Boneta, I., Soares, R., 2019. Chelonian use in Portugal: Evidence from Castelo Velho de Safara. Journal of Archaeological Science: Reports 28, 102054. 10.1016/j.jasrep.2019.102054

69. Nabais, M., Zilhão, J., 2019. The consumption of tortoise among Last Interglacial Iberian Neanderthals. Quaternary Science Reviews, Neanderthals: Ecology and Evolution 217, 225–246. 10.1016/j.quascirev.2019.03.024

70. Naksri, W., 2013. Origin and changes in continental turtle diversity from the Miocene to Holocene in Thailand (PhD thesis). Mahasarakham University, Maha Sarakham, Thailand.

71. Naksri, W., 2007. Appraisal of the evolution of testudinoid turtle diversity from the Oligocene and Neogene of Thailand (Master thesis). Mahasarakham University, Maha Sarakham, Thailand.

72. Naksri, W., Tong, H., Lauprasert, K., Suteethorn, V., Claude, J., 2013. A new species of *Cuora* (Testudines: Geoemydidae) from the Miocene of Thailand and its evolutionary significance. Geological Magazine 150, 908–922. 10.1017/S0016756812001082

73. Platt, S.G., Aung, S.H.N., Soe, M.M., Lwin, T., Platt, K., Walde, A.D., Rainwater, T.R., 2021. Predation on Translocated Burmese Star Tortoise (Geochelone platynota) by Asiatic Jackals (Canis aureus) and Wild Pigs (Sus scrofa) at a Wildlife Sanctuary in Myanmar. ccab 20, 133–138. 10.2744/CCB-1461.1

74. Pookajorn, S., 2001. New perspectives for Palaeolithic research in Thailand, in: Origine Des Peuplements et Chronologie Des Cultures Paléolithiques Dans Le Sud-Est Asiatique. Semenanjung, Paris, France, pp. 167–188.

75. Pritchard, P.C.H., Rabett, R.J., Piper, P.J., 2009. Distinguishing species of Geoemydid and Trionychid turtles from shell fragments: evidence from the Pleistocene at Niah Caves, Sarawak. Int. J. Osteoarchaeol. 19, 531–550. 10.1002/oa.1038

76. R Core Team, 2020. R: A language and environment for statistical computing.

77. Rabett, R.J., 2012. Tropical Subsistence Strategies at the End of the Last Glacial, in: Human Adaptation in the Asian Palaeolithic: Hominin Dispersal and Behaviour during the Late Quaternary. Cambridge University Press, Cambridge, UK, pp. 208–265.

78. Ramesh, M., 2008. Relative Abundance and Morphometrics of the Travancore Tortoise, Indotestudo travancorica, in the Indira Gandhi Wildlife Sanctuary, Southern Western Ghats, India. Chelonian Conservation and Biology 7, 108–113.

79. Rhodin, A.G.J., 1992. Chelonian Zooarchaeology of Eastern New England: Turtle bone remains from Cedar Swamp and other prehistoric sites. Bulletin of the Massachussetts Archaeological Society 53, 21– 30.

80. Rhodin, A.G.J., Iverson, J., Bour, R., Fritz, U., Georges, A., Shaffer, B.H., van Dijk, P.P., 2021. Turtles of the World: Annotated Checklist and Atlas of Taxonomy, Synonymy, Distribution, and Conservation Status (9th Ed.), Chelonian Research Monograph. Chelonian Research Foundation and Turtle Conservancy, United States of America.

81. Rouag, R., Benyacoub, S., Luiselli, L., El Mouden, E.H., Tiar, G., Ferrah, C., 2007. Population structure and demography of an Algerian population of the Moorish tortoise, Testudo graeca. Animal Biology 57, 267–279. 10.1163/157075607781753065

82. Santos, A., Mayor, P., Loureiro, L., Gilmore, M., Perez Peña, P., Bowler, M., Lemos, L., Svensson, M., Nekaris, K.A., Nijman, V., Valsecchi, J., Morcatty, T., 2020. Widespread Use of Traditional Techniques by Local People for Hunting the Yellow-Footed Tortoise (Chelonoidis denticulatus) Across the Amazon. Journal of Ethnobiology 40, 268. 10.2993/0278-0771- 40.2.268

83. Shoocongdej, R., 1996. Forager mobility organization in seasonal tropical environments: A view from Lang Kamnan Cave, western Thailand. (Thesis).

84. Som, S., Cottet, M., 2016. Rescue and relocation programme of turtles and tortoises and elongated tortoise monitoring programme in the Nam Theun 2 Reservoir (Laos). Hydroécol. Appl. 19, 383–406. 10.1051/hydro/2015007

85. Sophady, H., 2016. Archeo-stratigraphy of Laang Spean prehistoric site (Battambang province): a contribution to Cambodian prehistory (PhD thesis). Muséum national d’Histoire naturelle, Paris, France.

86. Sophady, H., Forestier, H., Zeitoun, V., Puaud, S., Frère, S., Celiberti, V., Westaway, K., Mourer, R., Mourer-Chauviré, C., Than, H., Billault, L., Tech, S., 2016. Laang Spean cave (Battambang province): A tale of occupation in Cambodia from the Late Upper Pleistocene to Holocene. Quaternary International, Southeast Asia: human evolution, dispersals and adaptation 416, 162–176. 10.1016/j.quaint.2015.07.049

87. Speth, J.D., Tchernov, E., 2002. Middle Paleolithic Tortoise Use at Kebara Cave (Israel). Journal of Archaeological Science 29, 471–483. 10.1006/jasc.2001.0740

88. Sriprateep, K., Aranyavalai, V., Aowphol, A., Thirakhupt, K., 2013. Population Structure and Reproduction of the Elongated Tortoise Indotestudo elongata (Blyth, 1853) at Ban Kok Village, Northeastern Thailand. Tropical Natural History 13, 21–37.

89. Stiner, M.C., Munro, N.D., Surovell, T.A., 2000. The Tortoise and the Hare: Small-Game Use, the Broad-Spectrum Revolution, and Paleolithic Demography. Current Anthropology 41, 39–79. 10.1086/300102

90. Taylor, E.H., 1970. The turtles and crocodiles of Thailand and adjacent waters; with a synoptic herpetological bibliography. The University of Kansas Science Bulletin 49, 87–179.

91. Theobald, W., 1868. Catalogue of the Reptiles of British Birma, embracing the Provinces of Pegu, Martaban, and Tenasserim; with descriptions of new or little-known species. Journal of the Linnean Society of London, Zoology 10, 4–67. 10.1111/j.1096- 3642.1868.tb02007.x

92. Thompson, J.C., Henshilwood, C.S., 2014. Tortoise taphonomy and tortoise butchery patterns at Blombos Cave, South Africa. Journal of Archaeological Science 41, 214–229. 10.1016/j.jas.2013.08.017

93. Treerayapiwat, C., 2005. Patterns of Habitation and Burial Activity in the Ban Rai Rock Shelter, Northwestern Thailand. Asian Perspectives 44, 231–245. 10.1353/asi.2005.0001

94. Tungittiplakornl, W., Dearden, P., 2002. Hunting and wildlife use in some Hmong communities in northern Thailand. Natural History Bulletin of the Siam Society 50, 57–73.

95. van Dijk, P.P., 1998. The Natural History of the Elongated Tortoise Indotestudo Elongata (Blyth, 1853) (Reptilia: Testudines) in a Hill Forest Mosaic in Western Thailand with Notes on Sympatric Turtle Species (Ph.D. dissertation). National University of Ireland, Galway, Ireland.

96. Van Vlack, H., 2014. Forager Subsistence Regimes in the Thai-Malay Peninsula: An Environmental Archaeological Case Study of Khao Toh Chong Rockshelter, Krabi (Master thesis. 4484). San José State University, San Jose.

97. Villa, P., Bon, F., Castel, J.-C., 2002. Fuel, fire and fireplaces in the Palaeolithic of western Europe. Review of Archaeology 23, 33–42.

98. Vu, T.L., 1994. The Hoabinhian animal complex. Vietnam Soc. Sci. 5, 70–74.

99. White, J.C., 2011. Emergence of cultural diversity in mainland Southeast Asia: A view from prehistory, in: Enfield, N.J. (Ed.), Dynamics of Human Diversity: The Case of Mainland Southeast Asia. Australian National University, Canberra, pp. 9–46.

100. Yen, E., 1977. Hoabinhian horticulture? The evidence and the questions from Northwest Thailand, in: Allen, J.G., Jones, R. (Eds.), Sunda and Sahul. Prehistoric Studies in Southeast Asia, Melanesia and Australia. Academic Press, London, pp. 567–599.

101. Zeitoun, V., Auetrakulvit, P., Zazzo, A., Pierret, A., Frère, S., Forestier, H., 2019. Discovery of an outstanding Hoabinhian site from the Late Pleistocene at Doi Pha Kan (Lampang province, northern Thailand). Archaeological Research in Asia 18, 1–16. 10.1016/j.ara.2019.01.002

102. Zeitoun, V., Forestier, H., Sophady, H., Puaud, S., Billault, L., 2012. Direct dating of a Neolithic burial in the Laang Spean cave (Battambang Province, Cambodia): First regional chrono-cultural implications. Comptes Rendus Palevol 11, 529–537. 10.1016/j.crpv.2012.06.006

103. Zeitoun, V., Forestier, H., Supaporn, N., 2008. Préhistoires au sud du Triangle d’Or, IRD Editions. ed. Paris, France.

104. Zeitoun, V., Heng, S., Forestier, H., 2021. The funeral cave of Laang Spean. Presented at the SEAMEO SPAFA International Conference on Southeast Asian Archaeology and Fine Arts, 13-17 December 2021, SEAMEO SPAFA, Bangkok, Thailand, pp. 1–14. 10.26721/spafa.pqcnu8815a-14

105. Znari, M., Germano, D.J., Macé, J.-C., 2005. Growth and population structure of the Moorish Tortoise (Testudo graeca graeca) in Westcentral Morocco: possible effects of over-collecting for the tourist trade. Journal of Arid Environments 62, 55–74. 10.1016/j.jaridenv.2004.11.013

106. Zuraina, M., 1994. The Excavation of Gua Gunung Runtuh and the Discovery of the Perak Man in Malaysia. Department of Museums and Antiquity Malaysia, Kuala Lumpur.

